# Discrete TrkB-expressing neurons of the dorsomedial hypothalamus regulate feeding and thermogenesis

**DOI:** 10.1101/2020.09.02.279745

**Authors:** Jessica Houtz, Guey-Ying Liao, Baoji Xu

**Affiliations:** Department of Neuroscience, The Scripps Research Institute Florida, Jupiter, FL 33458, USA

## Abstract

Mutations in the TrkB neurotrophin receptor lead to profound obesity in humans, and expression of TrkB in the dorsomedial hypothalamus (DMH) is critical for maintaining energy homeostasis. However, the functional implications of TrkB-expressing neurons in the DMH (DMH^TrkB^) on energy expenditure are unclear. Additionally, the neurocircuitry underlying the effect of DMH^TrkB^ neurons on energy homeostasis has not been explored. In this study, we show that activation of DMH^TrkB^ neurons leads to a robust increase in adaptive thermogenesis and energy expenditure without altering heart rate or blood pressure, while silencing DMH^TrkB^ neurons impairs thermogenesis. Furthermore, we reveal neuroanatomically and functionally distinct populations of DMH^TrkB^ neurons that regulate food intake or thermogenesis. Activation of DMH^TrkB^ neurons projecting to the raphe pallidus stimulates thermogenesis and increased energy expenditure, whereas DMH^TrkB^ neurons that send collaterals to the paraventricular hypothalamus and preoptic area inhibit feeding. Together, our findings provide evidence that DMH^TrkB^ neuronal activity plays an important role in regulating energy expenditure and delineate distinct neurocircuits that underly the separate effects of DMH^TrkB^ neuronal activity on food intake and thermogenesis.

**Brief summary:** This study shows that TrkB-expressing DMH neurons stimulate thermogenesis through projection to raphe pallidus, while inhibiting feeding through collaterals to paraventricular hypothalamus and preoptic area.

## Introduction

Impairments in energy homeostasis resulting from the compound effects of over-eating and sedentary lifestyles have led to a profound increase in the rate of obesity around the world, which has tripled in the past 50 years (1). Efforts to combat obesity have traditionally focused on lifestyle changes including diet and exercise; however, these strategies have met with limited success in sustaining weight loss (2, 3). Additionally, therapeutic strategies aimed at increasing energy expenditure or decreasing appetite have commonly failed due to adverse side effects on cardiovascular physiology (4–6). In order to achieve safe and sustained weight loss, it will be essential to understand the mechanisms that govern and coordinate discrete physiological processes that contribute to energy homeostasis.

The brain is a central regulator of energy balance which works through complex neural circuits that control feeding behavior and energy expenditure. Specifically, the dorsomedial hypothalamus (DMH) represents a highly connected and integral component of many neural circuits that contribute to energy balance including feeding, thermoregulation, metabolism, circadian rhythmicity, sleep, stress response, and reward seeking behavior (7–13). Adaptive thermogenesis is the process by which energy is converted into heat and occurs primarily in brown adipose tissue (BAT) in response to environmental cues. BAT has a particularly high capacity for dissipating energy from fat and thus represents an important component of energy homeostasis. The DMH is centrally positioned in an established thermoregulatory neurocircuit, receiving inputs from the preoptic area (POA) (8, 14, 15) and sending excitatory projections to pre-autonomic neurons in the raphe pallidus (RPa) (12, 16–18) that promote sympathetic activity in BAT leading to increased thermogenesis. Direct chemical stimulation of the DMH (19), or activation of select populations of thermogenic DMH neurons (8, 12, 17, 20) leads to increased body temperature and energy expenditure, but also significantly increases heart rate and blood pressure (12, 18, 20, 21). An inability to target increased sympathetic tone specifically in BAT without affecting other target tissues has greatly hampered strategies to treat obesity by targeting thermogenesis (5, 6).

In addition to its influence on energy expenditure, the DMH also represents an important brain region in the regulation of feeding (22–24). Lesioning studies support an orexigenic role for the DMH (22) which can promote food intake through inhibitory projections to either the paraventricular hypothalamus (PVH) (23) or the arcuate nucleus (ARC) (25). Despite these early findings, evidence has also emerged that demonstrates the importance of anorexigenic populations of DMH neurons (24, 26, 27). We previously established that the activity of DMH neurons expressing the neurotrophin receptor TrkB (DMH^TrkB^) is important for regulating feeding, showing that activation of DMH^TrkB^ neurons suppresses feeding and that deletion of the TrkB-encoding *Ntrk2* gene in the DMH results in hyperphagia and obesity (26). Furthermore, humans with mutations in the TrkB-encoding *NTRK2* gene exhibit severe obesity and impaired thermoregulation (28). However, it is unclear if activation of DMH^TrkB^ neurons has a direct influence on adaptive thermogenesis. Additionally, the neurocircuitry through which DMH^TrkB^ neurons govern feeding or energy expenditure are unknown.

Here, we demonstrate that DMH^TrkB^ neuronal activity potently promotes energy expenditure by elevating thermogenesis and physical activity with a notable lack of influence on heart rate and blood pressure. Additionally, we elucidate the neurocircuitry through which different populations of DMH^TrkB^ neurons influence discrete aspects of energy expenditure and food intake. We found that a novel DMH^TrkB^ →POA/PVH circuit, in which DMH^TrkB^ neurons send collaterals to both the PVH and the POA, regulates feeding and nutrient utilization, while RPa-projecting TrkB-expressing DMH (DMH^TrkB→RPa^) neurons promote energy expenditure via increased thermogenesis in BAT. These results show that DMH^TrkB^ neurons regulate feeding, physical activity and thermogenesis through distinct neural circuits without affecting the cardiovascular system.

## Results

### DMH^TrkB^ neurons are sensitive to environmental temperature

DMH neurons that respond to alterations in environmental temperature have been shown to promote thermogenesis (8, 29). In order to investigate a role for DMH^TrkB^ neurons in regulating thermogenesis, we first sought to determine if DMH^TrkB^ neurons are temperature sensitive. We crossed *Ntrk2*^*Cre-ER/+*^ mice (30) with the Cre-dependent tdTomato reporter mouse line (*Ai9*) (31) to generate *Ntrk2*^Cre-*ER/+*^;*Ai9* mice in which TrkB neurons are genetically labeled with tdTomato upon translocation of Cre-ERT2 protein to the nucleus induced by tamoxifen administration. Following labeling of TrkB neurons with tamoxifen, *Ntrk2*^*Cre-ER/+*^;*Ai9* mice were exposed to cold (10°C) or warm (39°C) temperatures, and immunostaining for Fos, a marker of activated neurons (32), was performed (Figure 1A-C). We found that exposure to cold temperature activated 10.4-11.8% of TrkB-expressing neurons throughout the anterior to posterior extent of the DMH (Bregma −1.46 – −2.18 mm), while warm temperature only activated 7.7-8.6% of TrkB neurons predominantly in the middle and posterior DMH (versus 1.9-3.1% activated DMH^TrkB^ neurons under the 30°C thermoneutral condition) (Figure 1D). Our results indicate that DMH^TrkB^ neurons respond to both increased and decreased ambient temperatures.

**Figure 1.**
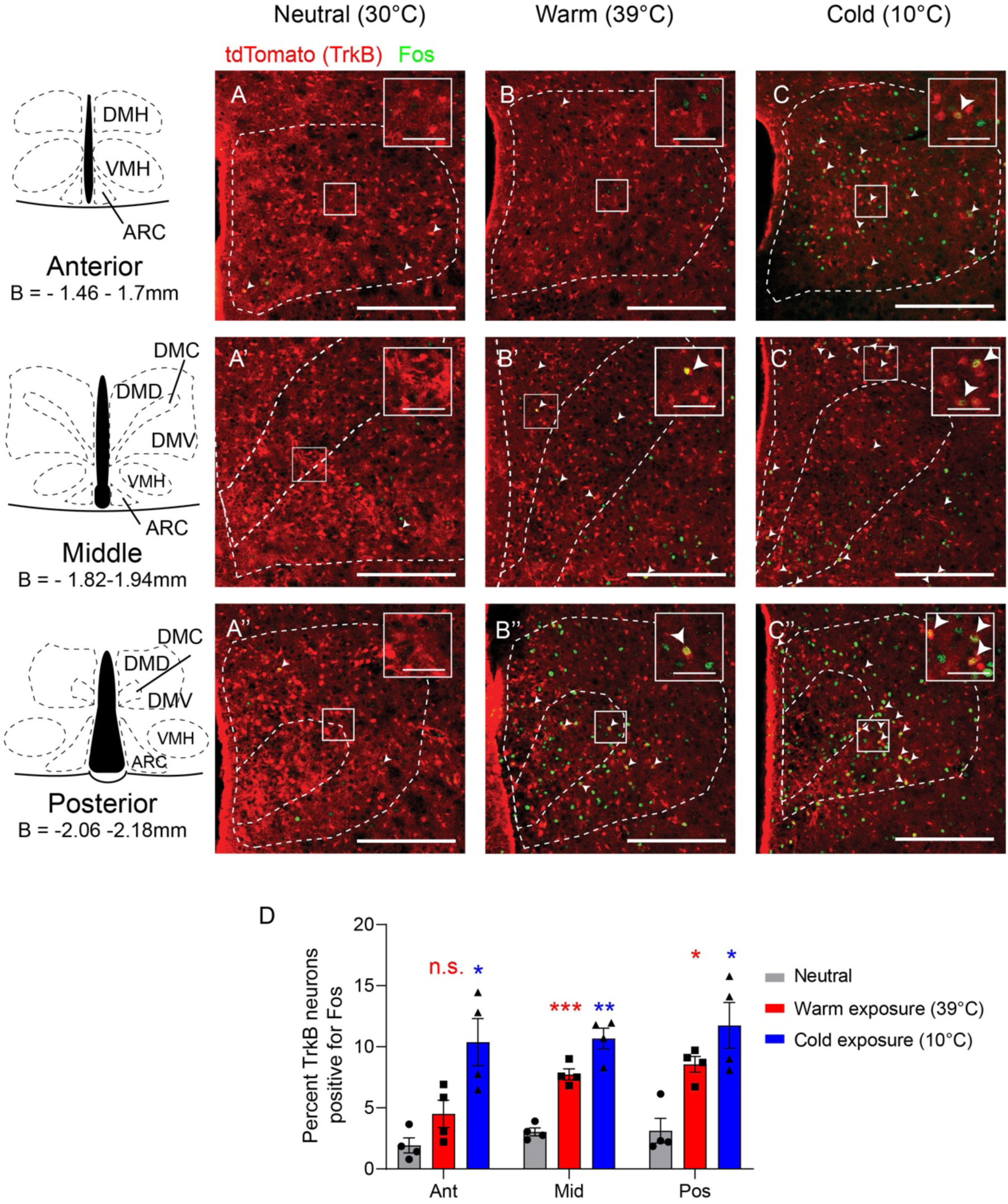
DMH^TrkB^ neurons are sensitive to changes in environmental temperature. Representative images of Fos staining (green) in the DMH of *Ntrk2*^*Cre-ER/+*^;*Ai9* reporter mice after exposure for 2 hours to (**A-A”**) thermoneutral (30°C), (**B-B”**) warm (39°C), or (**C-C”**) cold (10°C) temperatures. tdTomato (red) marks TrkB-expressing cells in reporter mice. Scale bars = 250 µm, inset scale bars = 50 µm (**D**) Quantification of Fos induction in TrkB-expressing neurons in the anterior (Ant), middle (Mid), and posterior (Pos) DMH after mice were exposed to different temperatures. 2-way ANOVA: temperature F_(2, 9)_ = 18.68, p=0.0006 *n* = 4 mice per condition; Dunnett’s post-test, n.s. = not significant, * p<0.05, ** p<0.01, *** p<0.001. Values represent mean ± s.e.m.. Abbreviations: DMH, dorsomedial hypothalamus; DMD, DMH dorsal division; DMC, DMH central division; DMV, DMH ventral division; VMH, ventromedial hypothalamus; ARC, arcuate nucleus; B, bregma.

### Chemogenetic manipulation of DMH^TrkB^ neurons alters thermogenesis and physical activity

The observed increase in Fos^+^ neurons in response to alterations in environmental temperature is consistent with a potential function for DMH^TrkB^ neurons in regulating body temperature through thermogenesis. In order to test the effect of manipulating DMH^TrkB^ neuronal activity on thermogenesis, we selectively expressed Designer Receptors Exclusively Activated by Designer Drugs (DREADD) in DMH^TrkB^ neurons (33). We delivered AAV virus expressing the excitatory DREADD, hM3Dq (AAV8-hSyn-DIO-hM3-mCherry) into the DMH of *Ntrk2*^*Cre-ER/+*^ mice to selectively activate DMH^TrkB^ neurons using the ligand clozapine N-oxide (CNO) (33). Mice expressing either control mCherry or hM3 in DMH^TrkB^ neurons (Figure 2A) were first acclimated to thermoneutrality (30°C) prior to measuring the effects of CNO administration on body temperature and energy expenditure in order to mitigate the influence of basal thermogenesis that occurs under normal housing temperatures (22°C) (34). Stimulation of DMH^TrkB^ neurons expressing hM3 with CNO (1.5 mg/kg) results in an acute and robust increase in body temperature compared with either vehicle stimulation in the same mice or CNO treatment in mice expressing control mCherry in DMH^TrkB^ neurons (Figure 2B, C). We also found that chemogenetic activation of DMH^TrkB^ neurons drives strong increases in both energy expenditure (Figure 2D, E) and locomotor activity (Figure 2H, I), while reducing the respiratory exchange ratio (RER) (VCO_2_/VO_2_) (Figure 2F, G) indicating a switch to the preferential utilization of fat as a metabolic substrate. Thermogenic activity in BAT drives both increased body temperature and energy expenditure as well as the utilization of fat, which results in a lower RER, thus, it is likely that DMH^TrkB^ neurons promote negative energy balance by regulating BAT activity. Many previous studies implicate DMH neuronal activity in promoting thermogenesis through an increase in sympathetic nerve activity (SNA) that also affects heart rate and blood pressure (12, 20, 21). Intriguingly, we found that stimulation of DMH^TrkB^ neurons in mice did not alter heart rate (Figure 2J) or mean arterial pressure (Figure 2K). Taken together, these findings support a selective role for DMH^TrkB^ neurons in regulating energy homeostasis by increasing fat metabolism, energy expenditure, and locomotor activity, without influencing cardiovascular physiology.

**Figure 2.**
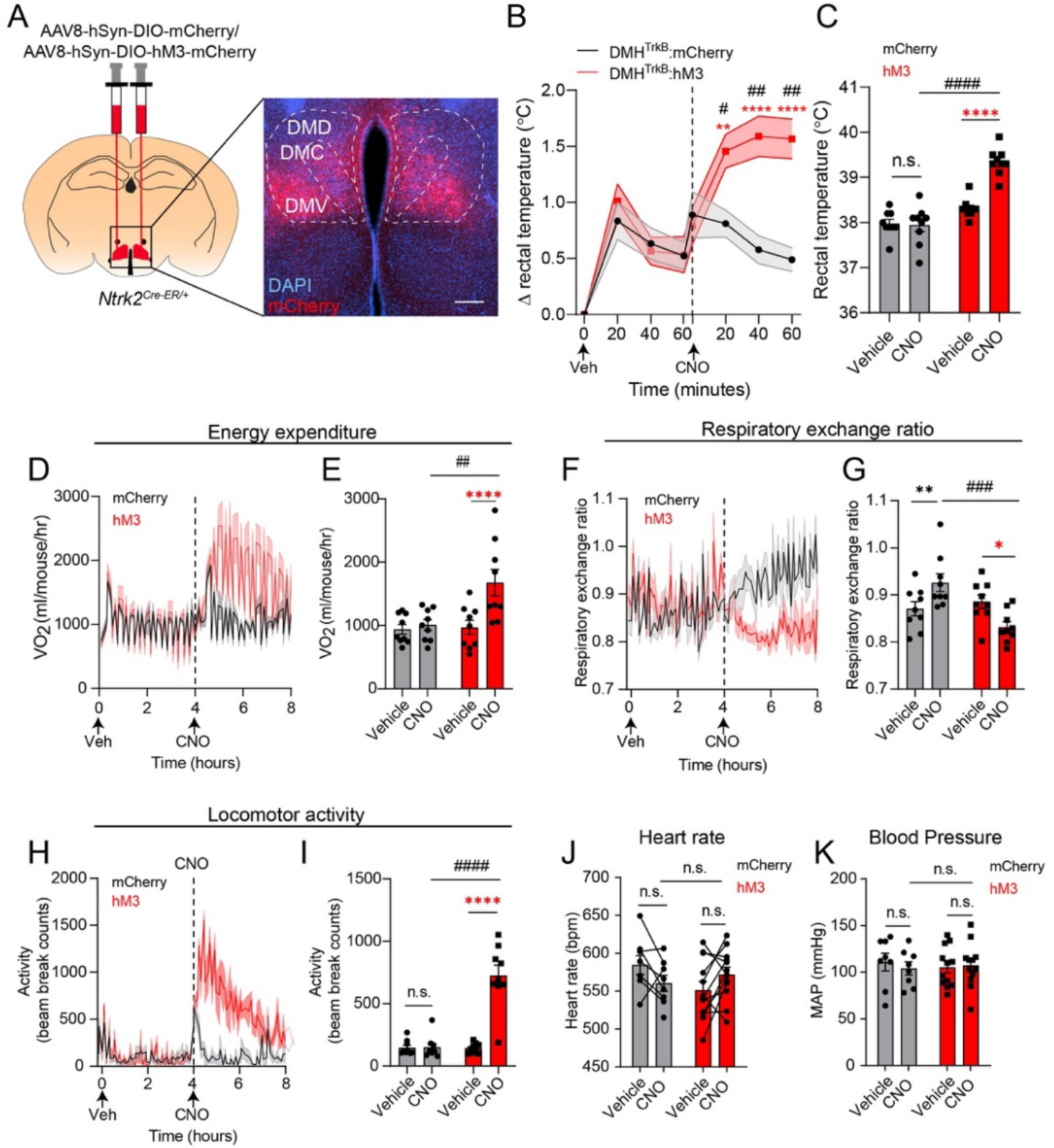
DMH^TrkB^ neuronal activity drives negative energy balance. (**A**) Schematic of bilateral stereotactic delivery of Cre-dependent AAV-expressing hM3-mCherry (AAV8-hSyn-DIO-hM3-mCherry) or mCherry (AAV8-hSyn-DIO-mCherry) into the DMH of *Ntrk2*^*Cre-ER/+*^ mice. Scale bar = 200 µm. (**B-I**) Mice housed at thermoneutrality expressing either mCherry (grey, *n* = 9), or hM3-mCherry (red, *n* = 9) in DMH^TrkB^ neurons were treated with vehicle (Veh) or CNO during the light cycle. (**B**) Rectal temperature. Two-way RM ANOVA: mCherry vs. hM3 F_(1, 16)_ = 10.11, p=0.0058. (**C**) Average rectal temperatures 60 minutes after treatment. Two-way RM ANOVA: mCherry vs hM3 F_(1, 16)_ = 36.55 p<0.0001. (**D**) Oxygen consumption (VO_2_) over 4 hours after treatment with Veh (0-4 hours) and CNO (4-8 hours). Mixed-effects model: mCherry *vs*. hM3 F_(1, 16)_ = 15.29, p=0.0012. (**E**) VO_2_ for the duration of the first four hours after treatment. Two-way RM ANOVA: mCherry vs. hM3 F_(1, 16)_ = 3.933, p=0.0648. (**F**) Respiratory exchange ratio over four hours following treatment. Mixed effects model: mCherry *vs*. hM3 expression F_(1, 16)_ = 16.65, p=0.0009. (**G**) Average respiratory exchange ratio for the duration of the first four hours after treatment. Two-way RM ANOVA: mCherry vs. hM3 expression F_(1, 16)_ = 5.072, p=0.0387. (**H**) Locomotor activity over 4 hours after treatment. Mixed effects model: mCherry vs. hM3 F_(1, 16)_ = 39.78, p<0.0001. (**I**) Average locomotor activity for the duration of the first four hours after treatment. Two-way RM ANOVA: mCherry vs. hM3 F_(1, 16)_ = 30.86, p<0.0001. (**J, K**) Heart rate and mean arterial pressure (MAP) in mice expressing mCherry or hM3 in DMH^TrkB^ neurons one hour after vehicle or CNO treatment. Two-way RM ANOVA for heart rate: effect of Veh vs CNO F_(1, 18)_ = 0.04648, p=0.8317; effect of mCherry vs. hM3 expression F_(1, 18)_ = 0.6447, p=0.4325. Two-way RM ANOVA for MAP: effect of Veh vs. CNO F_(1, 18)_ = 0.4427, p=0.5143; effect of mCherry vs. hM3 expression F_(1, 18)_ = 0. 01210, p=0.9136. *n* = 8 mCherry mice and 12 hM3 mice. Values represent mean ± s.e.m.. (**B-K**) Sidak post-test (red* hM3-Veh vs. hM3-CNO, black* mCherry-Veh vs. mCherry-CNO, # mCherry vs. hM3 post CNO: n.s. = not significant; *,# p<0.05; **.## p<0.01; ***,### p<0.001; ****,#### p<0.0001).

To further confirm a role for DMH^TrkB^ neurons in regulating energy expenditure by modulating BAT activity, we tested the necessity of DMH^TrkB^ neuronal activity for adaptive thermogenesis induced by cold exposure. We inactivated DMH^TrkB^ neurons by expressing the inhibitory DREADD receptor, hM4Di, in DMH^TrkB^ neurons (Figure 3A) and administering CNO (1.5 mg/kg). Silencing DMH^TrkB^ neurons in cold-housed (10°C) mice led to a reduction in body temperature (Figure 3B, C) and energy expenditure (Figure 3D, E). Additionally, we observed a significant increase in the RER (Figure 3F, G) which could be attributed either to an increase in feeding and subsequent availability of carbohydrate metabolic fuel that results from silencing DMH^TrkB^ neurons during the day (26), or the inhibition of lipolysis and shift to use of a carbohydrate fuel source that might be expected to occur concomitantly with the inhibition of BAT activity. Thus, we find that activation of DMH^TrkB^ neurons is necessary for the maintenance of body temperature in response to cold exposure that is achieved through increased energy expenditure.

**Figure 3.**
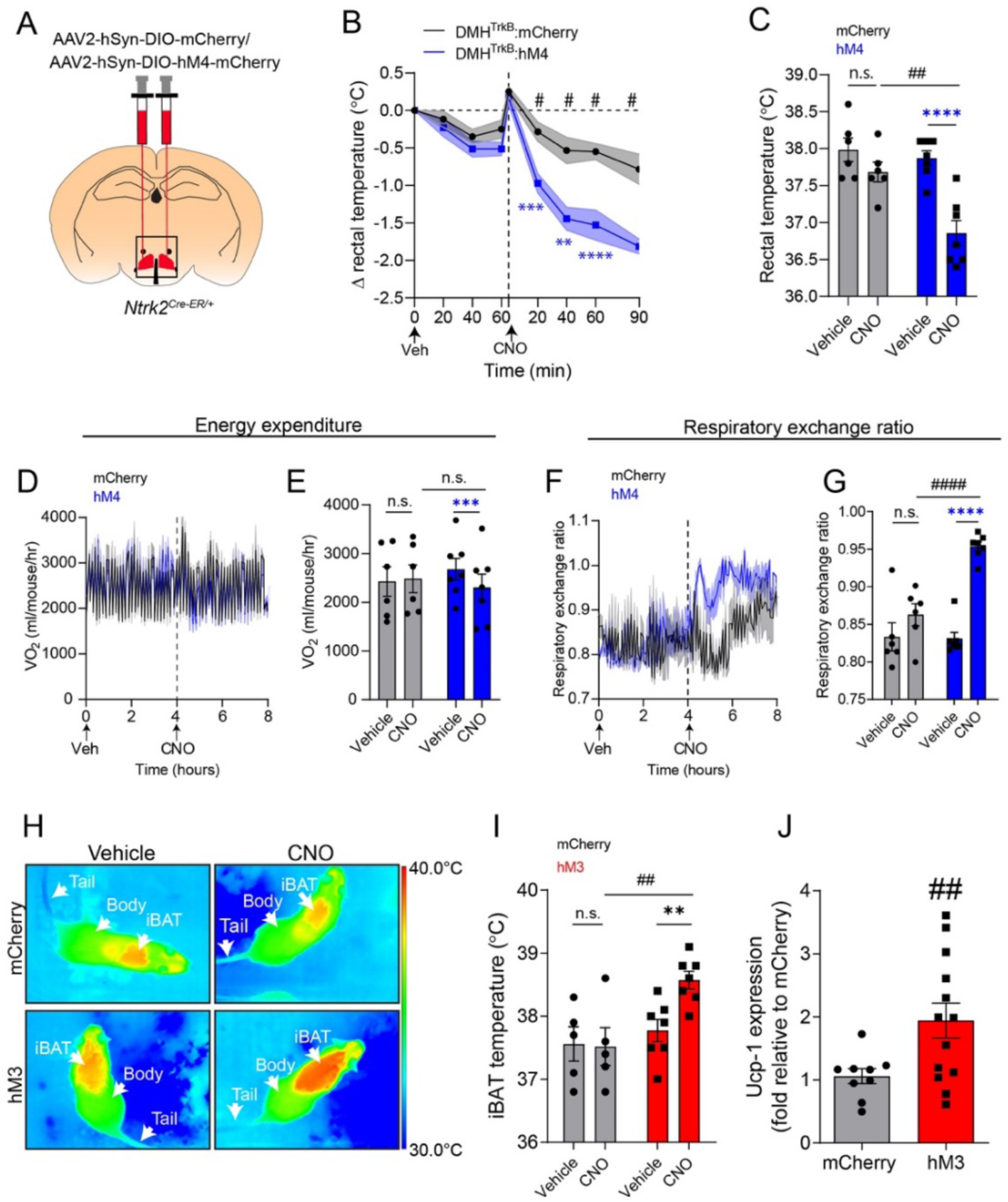
DMH^TrkB^ neurons are necessary for cold-induced thermogenesis. (**A**) Schematic of bilateral stereotactic delivery of Cre-dependent AAV-expressing inhibitory hM4-mCherry (AAV2-hSyn-DIO-hM4-mCherry) or mCherry (AAV2-hSyn-DIO-mCherry) into the DMH of *Ntrk2*^*Cre-ER/+*^. (**B-G**) Mice housed at 10°C expressing either mCherry (grey, *n* = 6) or hM4-mCherry (blue, *n* = 7) in DMH^TrkB^ neurons were treated with vehicle (Veh) or CNO during the light cycle. (**B**) Rectal temperature of mice following treatment with Veh (0-60 min) or CNO (0-90 min). 2-way RM ANOVA: mCherry vs. hM4 F_(1, 11)_ = 20.40, p=0.0009. (**C**) Average rectal temperatures 60 minutes after treatment. Two-way RM ANOVA: mCherry vs. hM4 F_(1, 11)_ = 7.577, p=0.0188. (**D**) Oxygen consumption over 4 hours after treatment. Mixed-effects model: mCherry vs. hM4 expression F_(1, 11)_ = 0.4110, p=0.5346. (**E**) Average VO_2_ for the duration of the first four hours after treatment. Two-way RM ANOVA: Veh vs. CNO F_(1, 11)_ = 12.96, p=0.0042. (**F**) Respiratory exchange ratio over 4 hours after treatment. Mixed-effects model: mCherry vs. hM4 F_(1, 11)_ = 39.28, p<0.0001. (**G**) Respiratory exchange ratio for the duration of the first four hours after treatment. Two-way RM ANOVA: mCherry vs. hM4 F_(1, 11)_ = 8.633, p=0.0135. (**H**) Representative thermal images and (**I**) iBAT temperature of mice housed at thermoneutrality expressing mCherry (*n* = 5) or hM3 (*n* = 7) in DMH^TrkB^ neurons 60 minutes post injection with either vehicle or CNO Two-way RM ANOVA: mCherry vs. hM3 F_(1, 10)_ = 6.350, p=0.0304. (**J**) Levels of *Ucp1* mRNA in iBAT from mice expressing hM3 or mCherry in DMH^TrkB^ neurons 2 hours following treatment with CNO (*n*= 9 mCherry, *n*= 13 hM3; unpaired, two-tailed t-test, p = 0.0207). Values represent mean ± s.e.m. (**B-K**) Sidak post-test (blue* hM4-Veh vs. hM4-CNO, black* mCherry-Veh vs. mCherry-CNO, # mCherry vs. hM4 post CNO: n.s. = not significant; *,# p<0.05; **.## p<0.01; ***,### p<0.001; ****,#### p<0.0001).

Due to the proximity of BAT to the surface of the skin, thermal imaging has emerged as a useful tool for evaluating thermogenesis (35). Therefore, we employed infrared thermography to further investigate the effect of stimulating DMH^TrkB^ neurons on promoting thermogenesis locally in BAT (Figure 3H). Following chemogenetic activation of DMH^TrkB^ neurons, we observed an increase in the subcutaneous temperature of the region above interscapular BAT (Figure 3I). Furthermore, we found an increase in the transcript level of the thermogenic marker, UCP-1 in BAT 2 hours after CNO stimulation in mice expressing the hM3 DREADD receptor in DMH^TrkB^ neurons but not mCherry (Figure 3J). In response to stress or cold stimulation, activation of the sympathetic nervous system elicits thermogenesis in BAT, but also leads to subcutaneous vasoconstriction to conserve body heat and results in a decrease in tail temperature in mice (15, 35, 36). However, we did not observe a change in tail temperature upon activation of DMH^TrkB^ neurons (Supplemental Figure 1A). The lack of vasoconstriction in response to activating DMH^TrkB^ neurons is consistent with our earlier observation that DMH^TrkB^ neurons do not appear to influence blood pressure. Together, this data supports a mechanism through which DMH^TkrB^ neurons elevate energy expenditure and body temperature by selectively activating thermogenesis in BAT.

The sympathetic nervous system is capable of acting in an organ specific manner, however the mechanisms governing this selectivity have not been defined (37). Since activation of DMH^TrkB^ neurons can induce thermogenesis without affecting heart rate, we hypothesized that these neurons may form a selective neurocircuit with sympathetic nerves that innervate BAT but not the heart. In order to test this possibility, we performed polysynaptic anterograde tracing in DMH^TrkB^ neurons using Cre-dependent HSV129ΔTK-tdTomato (Supplemental Figure 1B). Four days after infection, we observed connections between DMH^TrkB^ neurons and BAT as evident by dense tdTomato labeling in UCP-1 expressing BAT (Supplemental Figure 1C). In contrast, we did not see tdTomato labeling in either the heart or the sympathetic fibers innervating the heart (Supplemental Figure 1D). Our findings provide evidence that a DMH^TrkB^ neurons form selective polysynaptic connections with a specific population of sympathetic neurons that innervate BAT tissue.

### A DMH^TrkB^ → RPa neurocircuit regulates thermogenesis and energy expenditure, but not feeding

Given the multitude of ways that DMH^TrkB^ neuronal activity influences energy balance, through inhibition of feeding(26), stimulation of energy expenditure, and induction of physical activity (Figure 2), we next explored their efferent targets responsible for mediating these diverse processes. We performed anterograde tracing of DMH^TrkB^ neurons using AAV expressing Cre-dependent tdTomato (AAV2-CAG-FLEX-tdTomato) (Figure 4A). Following viral delivery to the DMH of *Ntrk2*^*Cre-ER/+*^ mice and induction with tamoxifen, DMH^TrkB^ neurons at the injection site were labeled with tdTomato (Figure 4B) and their projections were observed in multiple brain regions including the POA (Figure 4C), PVH (Figure 4D), ventrolateral periaqueductal grey matter (vlPAG) (Figure 4E), and RPa (Figure 4F). We also observed projections to the bed nucleus of the stria terminalis (BNST), anterior hypothalamus (AH), and ARC. Together, these data indicate that DMH^TrkB^ neurons have a broad projection field which may underly their functional diversity.

**Figure 4.**
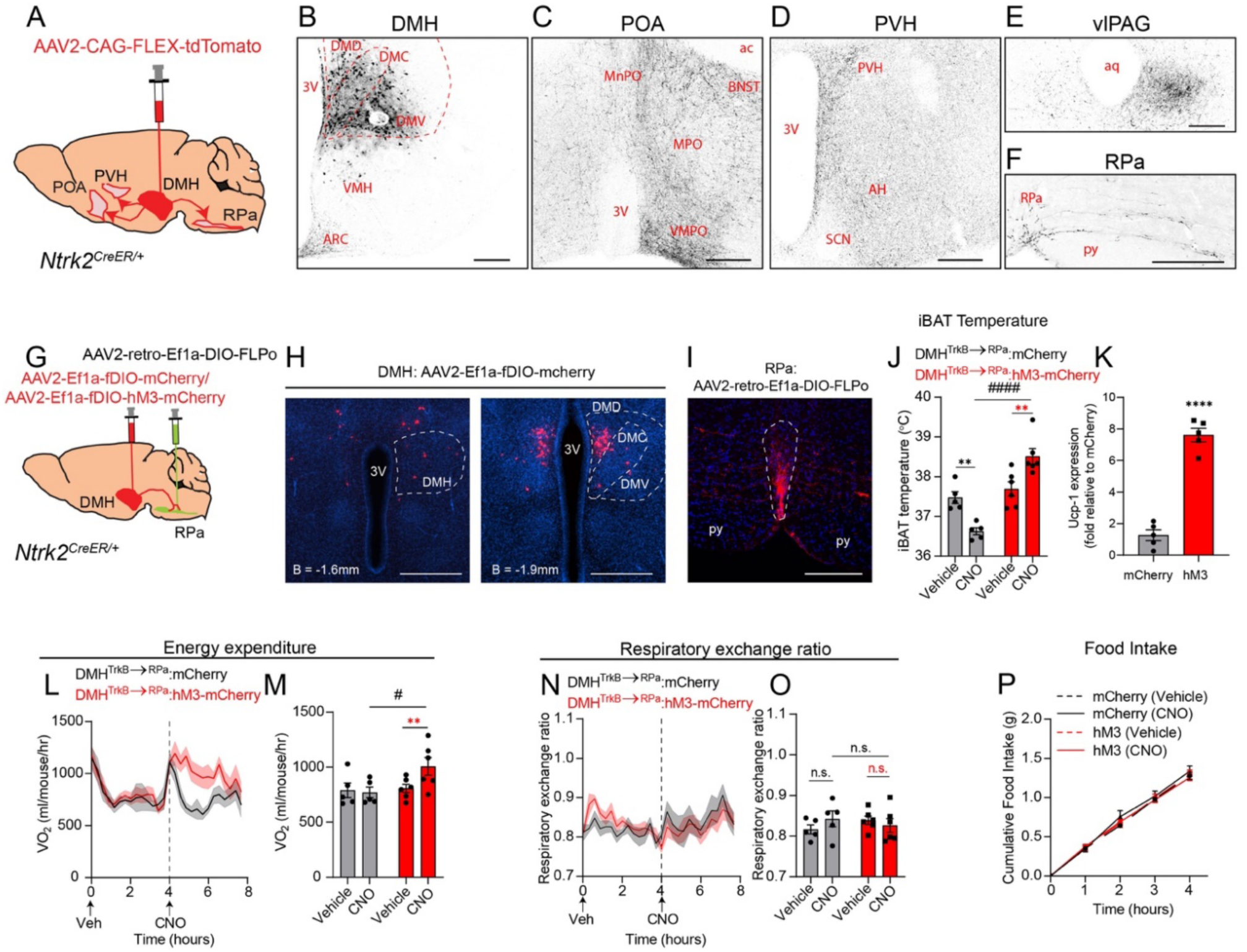
A DMH^TrkB^→RPa neurocircuit regulates energy expenditure and body temperature. (**A**) Diagram of anterograde tracing of DMH^TrkB^ neurons in *Ntrk2*^*Cre-ER/+*^ mice unilaterally injected with AAV2-CAG-FLEX-TdTomato. (**B**) Injection site. DMH, dorsomedial hypothalamus. (**C-F**) Projection targets: (**C**) preoptic area (POA) including the median preoptic area (MnPO), medial preoptic area (MPO), and ventral lateral preoptic area (VMPO) and the bed nucleus of the stria terminalis (BNST), (**D**) paraventricular hypothalamus (PVH) and anterior hypothalamus (AH), (**E**) ventrolateral part of periaqueductal grey (vlPAG), and (**F**) raphe pallidus (RPa). Scale bars = 200 µm. (**G**) Schematic of stereotactic delivery of retrograde AAV expressing Cre-dependent FLPo (AAV2-retro-Ef1a-DIO-FLPo) and AAV2-CMV-GFP to the RPa, and FLP-dependent (fDIO) mCherry or hM3-mCherry expressing virus to the DMH in *Ntrk2*^*Cre-ER/+*^ mice. (**H**) Expression of mCherry (red) in DMH^TrkB→RPa^ neurons. Scale bar = 500 µm. (**I**) mCherry-labeled DMH^TrkB→RPa^ terminals are detected in the RPa (outlined). Scale bar = 200 µm. (**J-O**) Mice housed at thermoneutrality and expressing hM3-mCherry (red, *n* = 6) or mCherry (grey, *n* = 5) in DMH^TrkB→RPa^ neurons were treated with vehicle (Veh) or CNO during the light cycle. (**J**) BAT temperature 60 minutes after Veh or CNO treatment. Two-way RM ANOVA: mCherry vs. hM3 F_(1, 9)_ = 29.24, p=0.0004. (**K**) Relative levels of *Ucp1* mRNA in BAT 2 hours post CNO treatment. Unpaired, two-tailed t-test, p<0.0001. (**L**) Oxygen consumption after treatment. Two-way RM ANOVA: mCherry vs. hM3 F_(1, 9)_ = 6.141, p=0.0351. (**M**) Average oxygen consumption for the duration of the first 4 hours after treatment. Two-way RM ANOVA: Veh vs. CNO F_(1, 9)_ = 7.013, p=0.0266. (**N**) Respiratory exchange ratio after treatment with vehicle or CNO. Two-way RM ANOVA: mCherry vs. hM3 F_(1, 9)_ = 0.2985, p=0.5985. (**O**) Average respiratory exchange ratio for the duration of the first 4 hours after treatment. Two-way RM ANOVA: mCherry vs. hM3 F_(1, 9)_ = 0.03257, p=0.8608. (**P**) Nocturnal food intake of mice expressing mCherry or hM3 in DMH^TrkB→RPa^ neurons after treatment with vehicle or CNO. Two-way RM ANOVA: mCherry vs. hM3 F_(1, 16)_ = 0.02664, p=0.8724. Values represent mean ± s.e.m. Sidak post-test (red* hM3-Veh vs. hM3-CNO, black* mCherry-Veh vs. mCherry-CNO, # mCherry vs. hM3 post CNO, n.s. = not significant, **.## p<0.01, #### p<0.0001). 3V-third ventricle, ac-anterior commissure, SCN-suprachiasmatic nucleus, py-pyramid

Sympathetic pre-motor neurons in the RPa are an established target of DMH neurons that are responsible for activating thermogenesis in BAT in response to cold or stress (12, 18, 38). Since we observed that DMH^TrkB^ neurons send projections to the RPa, we first reasoned that a DMH^TrkB^→RPa neurocircuit might be important for regulating thermogenesis and energy expenditure. To test this possibility, we developed a dual viral strategy to express the excitatory hM3 DREADD receptor in DMH^TrkB^ neurons in a projection-specific manner. We simultaneously delivered retrograde AAV expressing Cre-dependent FLPo recombinase (AAV2-retro-Ef1a-DIO-FLPo) and AAV2-CMV-GFP (to mark the injection site) to the RPa, and AAV expressing FLPo dependent hM3-mCherry or control mCherry (AAV2-Ef1a-fDIO-hM3-mCherry or AAV2-Ef1a-fDIO-mCherry) to the DMH in *Ntrk2*^*Cre-ER/+*^ mice (Figure 4G). TrkB-expressing neurons in the DMH that project to the RPa (DMH^TrkB→RPa^ neurons) become infected with AAV2-retro-EF1a-DIO-FLPo, and Cre expressed in TrkB neurons in *Ntrk2*^*Cre-ER/+*^ mice can induce expression of FLP recombinase allowing for subsequent expression of hM3-mCherry/mCherry. We confirmed expression of hM3-mCherry in neurons in the DMH (Figure 4H) and in axon terminals in the RPa (Figure 4I). When AAV2-Ef1a-fDIO-hM3-mCherry or AAV2-Ef1a-fDIO-mCherry were injected into the DMH simultaneously with AAV2-retro-Ef1a-DIO-FLPo into the RPa of wildtype mice that lack Cre expression in TrkB-expressing neurons, we saw minimal signal in the red channel that was likely due to cell debris (Supplemental Figure 2A-C). Additionally, injection of AAV2-Ef1a-fDIO-hM3-mCherry or AAV2-Ef1a-fDIO-mCherry alone into the DMH of *Ntrk2*^*Cre-ER/+*^ mice without a second injection of AAV2-retro-Ef1a-DIO-FLPo into the RPa was insufficient to induce expression of mCherry or hM3-mCherry (Supplemental Figure 2D-F). As with injections into wildtype mice, red fluorescence observed in *Ntrk2*^*Cre- ER/+*^ mice injected with AAV2-Ef1a-fDIO-hM3-mCherry is likely due to cellular debris or glial scar formation at the injection site since no signal is apparent in cell processes. Treatment with CNO did not result in increased Fos expression in DMH^TrkB→RPa^ neurons expressing mCherry compared with hM3 (Supplemental Figure 2G, H). These results support our ability to target specific DMH^TrkB^ neurons for chemogenetic activation based on their projection target.

We then tested the effect of activating the DMH^TrkB^→RPa neurocircuit on thermogenesis in BAT and energy expenditure. Treatment with CNO, but not vehicle, increased BAT temperature (Figure 4J) and expression of a thermogenic marker, *Ucp1*, (Figure 4K) in *Ntrk2*^*Cre-ER/+*^ mice expressing hM3 in DMH^TrkB→RPa^ neurons compared with mCherry expressing control mice. Likewise, we observed an increase in oxygen consumption in response to activating DMH^TrkB→RPa^ neurons (Figure 4L, M). In contrast to the effect of activating all DMH^TrkB^ neurons, stimulation of DMH^TrkB→RPa^ neurons did not alter RER (Figure 4N, O) or physical activity (Supplemental Figure 3A, B). We previously found that activation of DMH^TrkB^ neurons suppresses nocturnal feeding (26), however the specific stimulation of a DMH^TrkB^→RPa circuit has no effect on food intake (Figure 4P). Although activation of DMH^TrkB→RPa^ neurons increased thermogenesis in BAT to a similar level as previously observed (DMH^TrkB^: hM3 at 60 min post-CNO = 38.6±0.1°C vs. DMH^TrkB→RPa^: hM3 at 60 min post-CNO = 38.5±0.2°C), the DMH^TrkB^→RPa neurocircuit was not sufficient to drive total energy expenditure to the same extent or duration as that observed for activation of all DMH^TrkB^ neurons (DMH^TrkB^:hM3, 73% vs. DMH^TrkB→RPa^:hM3, 25% increase in VO_2_ for CNO over vehicle). Thus, we find that a DMH^TrkB^→RPa neurocircuit contributes to the net negative effect of DMH^TrkB^ neurons on energy balance by increasing energy expenditure and thermogenesis but is not involved in the regulation of fat metabolism or feeding.

### DMH^TrkB^ neurons send collaterals to the PVH and POA to regulate feeding and substrate utilization

We observed that activation of DMH^TrkB^ neurons drives high levels of oxygen consumption while at the same time directing preferential utilization of fat as a metabolic fuel source (Figure 2) and inhibiting homeostatic feeding (26). Since DMH^TrkB→RPa^ neurons do not influence RER or feeding, we aimed to determine the identity of a DMH^TrkB^ neurocircuit that participates in these other components of energy balance. Neurons in the PVH that express BDNF (a high affinity TrkB ligand) have been implicated in regulating food intake, energy expenditure, and locomotor activity, however the identity of their presynaptic partners is unknown (39). Our anterograde tracing results indicate that the PVH receives projections from DMH^TrkB^ neurons (Figure 4D), thus we hypothesized that a DMH^TrkB^→PVH pathway might account for the effect of DMH^TrkB^ neuronal activity on feeding. In order to test this possibility, we used the same projection-specific viral strategy described earlier to express hM3-mCherry or mCherry in PVH-projecting DMH^TrkB^ neurons (Figure 5A, B). DMH^TrkB→PVH^ neurons expressing mCherry are present in the middle-to-posterior ventral DMH (DMV) and the medial-ventral part of the central DMH (DMC) (Figure 5B). We confirmed proper targeting of the PVH by co-injection of AAV-GFP with AAV-retro-DIO-FLPo. PVH neurons at the injection site expressed GFP and received dense innervation by mCherry^+^ DMH^TrkB→PVH^ fibers (Figure 5C, C’). These results indicate successful targeting of a DMH^TrkB^→PVH neurocircuit for chemogenetic manipulation and reveal that DMH^TrkB→PVH^ neurons are spatially distinct from DMH^TrkB→RPa^ neurons.

**Figure 5.**
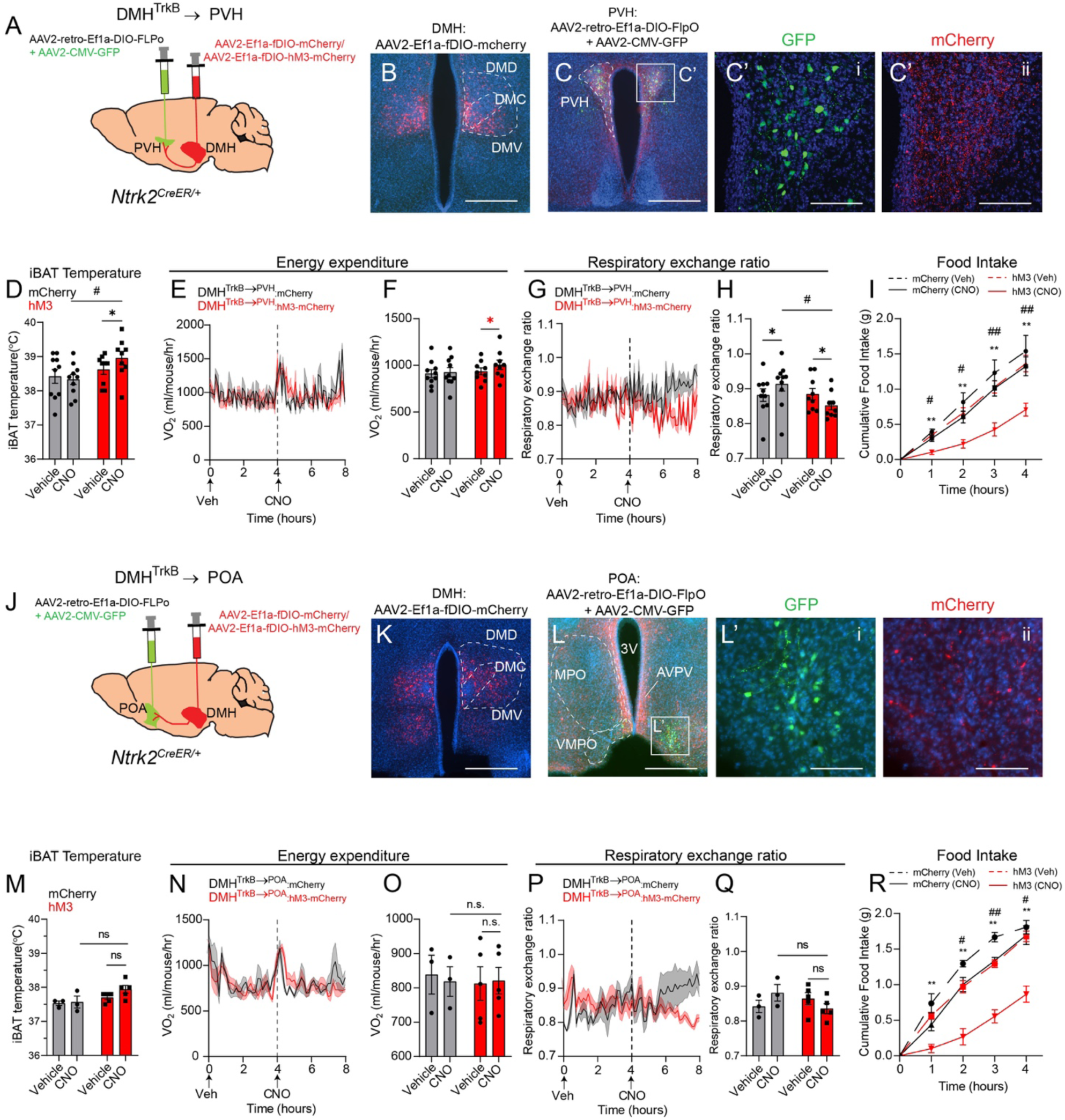
DMH^TrkB^ neurons project to the PVH and POA to regulate feeding and metabolism. **(A, J)** Strategy for projection-specific targeting of mCherry or hM3-mCherry expression in DMH^TrkB^ neurons. Retrograde AAV expressing Cre-dependent FLPo (AAV2-retro-Ef1a-DIO-FLPo) and AAV-GFP are delivered to the PVH (**A**) or POA (**J**), while Flp-dependent mCherry or hM3-mCherry expressing virus is injected into the DMH in *Ntrk2*^*Cre-ER/+*^ mice. (**B**) Expression of mCherry in DMH^TrkB→PVH^ neurons. Scale bar, 500 µm. (**C, C’**) Expression of GFP marking the injection site of retrograde AAV2-retro-Ef1a-DIO-FLPo in the PVH (**C’i**) and mCherry in axonal terminals of DMH^TrkB^ neurons in the PVH (**C’ii**). **(B, C)** Scale bars represent 500 µm in (**B**) and 100 µm in (**C’**). (**D**-**H**) Mice housed at thermoneutrality and expressing mCherry (grey, *n* = 10) or hM3-mCherry (red, *n* = 9) in DMH^TrkB→PVH^ neurons were treated with vehicle (Veh) and then CNO during the light cycle. (**D**) iBAT temperature. Two-way RM ANOVA: mCherry vs. hM3 F_(1, 17)_ = 3.178, p=0.0925. (**E**) Oxygen consumption. Mixed-effects model: mCherry vs. hM3 F_(1, 17)_ = 0.008586, p=0.0002. (**F**) Average oxygen consumption for the duration of 4 hours following treatment. Two-way RM ANOVA: Veh vs. CNO F_(1, 17)_ = 7.704, p=0.0130. (**G**) Respiratory exchange ratio. Mixed-effects model: mCherry vs. hM3 F_(1, 17)_ = 5.311, p=0.0341. (**H**) Average respiratory exchange ratio for the duration of 4 hours following treatment. Two-way RM ANOVA: mCherry vs. hM3 F_(1, 17)_ = 1.614, p=0.008168. **(I)** Nocturnal food intake of mice expressing mCherry or hM3 in DMH^TrkB→PVH^ neurons post injection of vehicle or CNO. Two-way RM ANOVA: mCherry vs. hM3 F_(1, 17)_ = 4.913, p=0.0406. (**K**) Expression of mCherry in DMH^TrkB→POA^ neurons. Scale bar = 500 µm (**L, L’**) Expression of GFP marking the injection site of AAV2-retro-Ef1a-DIO-FLPo in the POA and mCherry in axonal terminals of DMH^TrkB^ neurons in the POA. Scale bars = 500 µm in (**L**) and 100 µm (**L’**). (**M**-**O**) Mice housed at thermoneutrality and expressing mCherry (grey, *n* = 3) or hM3-mCherry (red, *n* = 5) in DMH^TrkB→POA^ neurons were treated with Veh and then CNO during the light cycle. (**M**) iBAT temperature. Two-way RM ANOVA: mCherry vs. hM3 F_(1, 6)_ = 5.031, p=0.0661. (**N**) Oxygen consumption. Two-way RM ANOVA: mCherry vs. hM3 F_(1, 6)_ = 0.0002876, p=0.9870. (**O**) Average oxygen consumption for the duration of 4 hours following treatment. Two-way RM ANOVA; mCherry vs. hM3 F_(1, 6)_ = 0.03586, p=0.8560). (**P**) Respiratory exchange ratio. Two-way RM ANOVA: mCherry vs. hM3 F_(1, 6)_ = 2.985, p=0.1348. (**Q**) Average respiratory ratio for the duration of 4 hours following treatment. Two-way RM ANOVA; mCherry vs. CNO F_(1, 6)_ = 0.2390, p=0.6423. **(R)** Nocturnal food intake of mice expressing mCherry or hM3 in DMH^TrkB→POA^ neurons. Two-way RM ANOVA: mCherry vs. hM3 F_(1, 6)_ = 24.98, p=0.0025. Values represent mean ± s.e.m. Sidak post-test (*Veh vs. CNO, #mCherry vs. hM3; *,#p<0.05 **,## p<0.01).

Similar to the effect of activating the DMH^TrkB^→RPa circuit, activation of DMH^TrkB→PVH^ neurons increases BAT temperature one hour after treatment with CNO (Figure 5D). Despite this evidence indicating a potential role for DMH^TrkB→PVH^ neurons in thermogenesis, activating this circuit did not have a large effect on energy expenditure. Following stimulation with CNO, mice expressing hM3 in DMH^TrkB→PVH^ neurons showed a slight increase in oxygen consumption (Figure 5E, F) which was restricted to the first hour following stimulation (Figure 5E). As was observed for activation of DMH^TrkB→RPa^ neurons, the DMH^TrkB^→PVH neurocircuit was not sufficient to drive oxygen consumption to the same extent as activation of all DMH^TrkB^ neurons (DMH^TrkB^:hM3, 73% vs. DMH^TrkB→PVH^:hM3, 8% increase in VO_2_ for CNO over vehicle), nor did it have any influence on physical activity (Supplemental Figure 3C, D). However, stimulation of the DMH^TrkB^→PVH circuit lowers the RER (Figure 5G, H) and inhibits nocturnal feeding in mice (Figure 5I). Thus, DMH^TrkB→PVH^ neurons contribute to negative energy balance predominantly by regulating nutrient consumption and utilization.

Another population of BDNF-expressing neurons, located in the POA (POA^BDNF^), has also been described that project to the DMH and influence body temperature by regulating BAT activity (15). Since DMH^TrkB^ neurons project to the POA, we were curious if DMH^TrkB→POA^ neurons could feed back to inhibit these POA^BDNF^ neurons in order to drive thermogenesis and increase body temperature. Thus, we targeted expression of hM3-mCherry or mCherry to DMH^TrkB→POA^ neurons (Figure 5J, K). Expression of GFP at the injection site was predominantly restricted to the ventromedial preoptic area (VMPO) (Figure 5L, L’), where warm-sensitive BDNF-expressing neurons have previously been reported to be concentrated (15). While mCherry^+^ DMH^TrkB→POA^ axon terminals were evident in the VMPO (Figure 5L, L’ii), we also noted collateralization to the MPO as well as the anteroventral periventricular nucleus (AVPV) (Figure 5L). This data indicates that DMH^TrkB→POA^ neurons have a large projection field within the POA which is likely not restricted to POA^BDNF^ neurons. Activation of DMH^TrkB→POA^ neurons expressing hM3-mCherry with CNO had no significant influence on BAT temperature (Figure 5M), oxygen consumption (Figure 5N, O), RER (Figure 5P, Q) or physical activity (Supplemental Figure 3E, F). However, we noted a slight trend toward a lower RER in the 4^th^ hour following CNO treatment in mice expressing hM3-mCherry compared with mCherry (Figure 5P). As was observed for the activation of the DMH^TrkB^→PVH circuit, we saw that treatment with CNO inhibited nocturnal feeding in mice expressing hM3 compared with mCherry expression in DMH^TrkB→POA^ neurons or compared with vehicle treatment (Figure 5R). Together, these results demonstrate that DMH^TrkB→POA^ and DMH^TrkB→PVH^ neurons have a redundant function in the regulation of food intake.

In addition to their functional redundancy, we found that DMH^TrkB→PVH^ and DMH^TrkB→POA^ neurons have a similar distribution within the DMH (Figure 5B, K). These results raise the possibility that a single population of DMH^TrkB^ neurons might send projections to both the PVH and the POA. Indeed, we observed that DMH^TrkB→PVH^ neurons send collaterals to the POA, including both the MPO and VMPO, and the BNST (Supplemental Figure 4C, C’), but not the RPa (Supplemental Figure 4D). Similarly, we observed DMH^TrkB→POA^ collaterals in the PVH (Supplemental Figure 4G, G’), but DMH^TrkB→RPa^ collaterals were not apparent in either the POA (Supplemental Figure 4J) or the PVH (Supplemental Figure 4K). Thus, we find that there are at least two populations of TrkB neurons in the DMH that either project to the RPa or send collaterals to the PVH and POA.

In order to further confirm the existence of DMH^TrkB^ neurons that collateralize to both the PVH and the POA, we simultaneously injected CTB labeled with two different fluorophores (Alexa-488 or -647) into the PVH and the POA of *Ntrk2*^*Cre-ER/+*^;*Ai9/+* mice in which labeling of TrkB neurons with tdTomato was previously induced with tamoxifen (Supplemental Figure 4L, O). Neurons in the DMH that project to the PVH were evident by labeling with Alexa-488, while those that project to the POA were labeled with Alexa-647 (Supplemental Figure 3O), and TrkB expressing neurons were labeled with tdTomato (Supplemental Figure 4P). PVH- and POA-projecting DMH neurons were concentrated in the ventral part of the middle-to-posterior DMH. Upon further inspection, we found that most DMH neurons that send collateral projections to both the POA and PVH express TrkB (Supplemental Figure 4Q-T, arrowheads). These findings support the existence of a specific population of DMH^TrkB^ neurons that send projections to both the POA and the PVH.

## Discussion

Expression of TrkB in the brain is essential for maintaining normal body weight in both humans and rodents (28, 40, 41). We have previously reported that TrkB expressing neurons in the DMH (26) and the PVH (42) are essential for regulating food intake, and DMH^TrkB^ neuronal activity modulates homeostatic feeding. Although we observed that loss of TrkB signaling in the DMH also leads to reduced oxygen consumption and physical activity, these consequences could be secondary to excessive weight gain. In the present study, we establish that DMH^TrkB^ neuronal activity has a direct influence on energy expenditure and body temperature that does not extend to influence components of cardiovascular physiology which are typically affected by other DMH neuron populations. Furthermore, we reveal diverging DMH^TrkB^ neurocircuits that are responsible for regulating adaptive thermogenesis, physical activity, fat metabolism, and appetite. Together, our findings support an integral role for DMH^TrkB^ in coordinating multiple arms of energy balance and provide critical insight into the neural mechanisms underlying their functions.

The DMH is recognized as a brain center that predominantly promotes energy expenditure in part through the activation of thermogenesis in BAT (19, 43). Neurons located in the anterior and dorsal compartments of the DMH are sensitive to temperature reductions and project to the RPa to mediate increased sympathetic tone in BAT leading to thermogenesis (12, 20, 44). Consistent with these reports, we found that cold exposure increased Fos expression in DMH^TrkB^ neurons, and that activation of DMH^TrkB→RPa^ neurons localized in the anterior and dorsal DMH induces heat production and *Ucp1* expression in BAT. DMH projections to pre-autonomic neurons in the RPa act to increase sympathetic tone which has been demonstrated to promote tachycardia and hypertension in addition to thermogenesis (12, 20, 21). Surprisingly, we observed that activation of DMH^TrkB^ neurons has no obvious effect on cardiovascular physiology as evident both by unchanged measurements of heart rate and blood pressure as well as tail temperature, an indicator of vasodilation or vasoconstriction. Our anterograde neural tracing results suggest that this specificity may be attributed to the selective targeting of neural circuits connecting to BAT, but not other target tissues of the SNS, namely the heart. Separate populations of sympathetic neurons have been shown to innervate either white adipose tissue (WAT) or BAT, indicating selectivity in sympathetic output at the level of the peripheral nervous system (37). Although DMH^TrkB^ neurons might form polysynaptic connections with specific sympathetic neurons that regulate BAT activity, it is unclear how this selectivity is retained at the level of the RPa. Thus, future investigations will be essential for determining the molecular diversity of neurons in the RPa that form connections with different populations of thermogenic DMH neurons.

Given that activation of DMH^TrkB→RPa^ neurons does not drive energy expenditure to the full extent seen by stimulating the global DMH^TrkB^ population, other DMH^TrkB^ neurocircuits likely contribute to increased total energy expenditure. DMH^TrkB→PVH^ neurons might help facilitate BAT activity by inhibiting NTS-projecting PVH neurons that negatively regulate the RPa (45, 46) or by exciting PVH^BDNF^ neurons that are polysynaptically connected to BAT (39). This possibility is in line with the subtle thermogenic effect of activating a DMH^TrkB^→PVH neurocircuit. DMH^TrkB→PVH^ neurons could also augment thermogenesis through neuroendocrine signaling (47, 48). However, DMH^TrkB^ neuron activation does not alter heart rate and blood pressure, while many PVH-derived factors, such as cortocotropin-releasing peptide (CRH) and thyroid-releasing hormone (TRH) have well established influences on cardiovascular physiology.

Physical activity represents another significant component of total energy expenditure. In addition to the intrinsic energy demands of increased physical activity, exercise is known to stimulate the production of thermogenic factors such as fibroblast growth factor-21 (FGF-21) and irisin which could also contribute to thermogenesis related energy expenditure (49). None of DMH^TrkB^→RPa, DMH^TrkB^→PVH, or DMH^TrkB^→POA neurocircuits elicits increased physical activity upon activation, thus an undefined DMH^TrkB^ activity-promoting neurocircuit may contribute to the collective increase in energy expenditure seen by activating DMH^TrkB^ neurons. Recent studies have revealed an important role for neurons in the BNST in promoting arousal and wakefulness associated with increased locomotor activity (50–52). Furthermore, projections from calretinin neurons in the paraventricular thalamus can mediate an increase in activity associated with starvation (52). Although we do not investigate the effect of DMH^TrkB^ activity on sleep architecture, we see that activation of all DMH^TrkB^ neurons robustly triggers increased locomotor activity and send projections to the BNST (Figure 4C). DMH^TrkB→PVH/POA^ neurons appear to also send collaterals to the BNST (Supplemental Figure 5C), however activation of these neurocircuits did not alter locomotor activity. The BNST is a large brain structure that contains many subregions and diverse neuron populations (50). Thus, it is possible that DMH^TrkB^→BNST projections and collaterals in the BNST from DMH^TrkB→PVH/POA^ neurons represent non-overlapping neurocircuits that target different populations of neurons in the BNST. A DMH^TrkB^→BNST neurocircuit could therefore represent a pathway through which locomotor activity is appropriately coordinated and integrated with other processes that mediate energy homeostasis.

Food intake represents the positive arm of energy balance which shares equal import with energy expenditure for regulating body weight. Our previous studies indicate that TrkB expression in the DMH and neuronal activity of DMH^TrkB^ neurons are necessary for the control of homeostatic feeding. Since activation of DMH^TrkB→RPa^ neurons does not affect nocturnal feeding, we surmised that a different DMH^TrkB^ neurocircuit might regulate feeding. Neurons in the DMH, including DMH^TrkB^ neurons, send dense projections to the PVH which is comprised of several anorexogenic glutamatergic populations, including those that express BDNF (39, 53, 54). BDNF signaling through TrkB is critical for neural circuit development and synaptic plasticity (55), thus PVH^BDNF^ neurons represent a potential neural substrate for DMH^TrkB→PVH^ neurons that govern feeding. BDNF secretion from postsynaptic neurons in the PVH could presynaptically modulate DMH^TrkB^ neuronal activity (56, 57). This effect might be expected to yield prolonged alteration in signaling that could contribute to homeostatic feeding as compared with the relatively short-term pre-prandial anorexic effects of previously described DMH^LepR^→ARC^AgRP^ neurocircuit (24). Indeed, application of BDNF increases firing of PVH-TrkB neurons after a delay of up to 40 minutes, consistent with the relatively slow kinetics of presynaptic components of potentiation in the hippocampus (58).

DMH^TrkB→PVH^ neurons collateralize extensively to the POA which also contains BDNF-expressing neurons. However, stimulation of POA^BDNF^ neurons has a potent hypothermic effect on body temperature and drastically reduces energy expenditure resulting in a hibernation-like state (15, 36), and are therefore an unlikely target of DMH^TrkB^ neurons. Despite significant overlap in retrogradely labeled DMH^TrkB^ populations that project to the PVH and the POA, activation of DMH^TrkB→POA^ neurons was not sufficient to significantly increase BAT temperature or energy expenditure as was observed for activation of DMH^TrkB→PVH^ neurons. This discrepancy may be due to an incomplete activation of all DMH^TrkB→POA^ neurons, attributed to the limited expression of retrograde FLPo-expressing virus at the VMPO. Alternatively, DMH^TrkB→PVH^ neurons that drive thermogenesis and energy expenditure could represent a sub-population that does not send collaterals to the POA. Nevertheless, we find that activation of a DMH^TrkB^→POA neurocircuit has a potent inhibitory effect on feeding.

The POA is recognized as a critical brain region in the homeostatic regulation of body temperature, sleep, osmolarity, and reproduction, however the effect of the POA on food intake is largely unexplored (8, 59–61). Under warm ambient temperatures, the metabolic demand for thermogenesis is reduced and thus energy expenditure and food intake are appropriately lowered to maintain body weight homeostasis (34, 62). Leptin receptor expressing neurons in the POA that drive hypothermia have been implicated in temperature-dependent regulation of body weight in part by inhibiting feeding (44) and recent studies have revealed a large degree of diversity in known populations of POA neurons (63, 64). In future experiments, it will be interesting to determine if DMH^TrkB→POA^ neurons target a previously unrecognized subpopulation of POA neurons responsible for regulating food intake without influencing body temperature or energy expenditure.

Activation of POA neurons has a well-established effect of lowering body temperature (8, 15, 44). Recent investigations have revealed that hypothermia inducing POA neurons may in fact be regulating entry into a hibernation-like state (36) or daily torpor (64) in which energy expenditure is reduced in parallel with body temperature. During these states of reduced energy expenditure and activity, mice also show a reduction in RER characteristic of a switch to lipid utilization as an energy source. Daily variation in RER, with lower RER occurring during the day when mice are not eating and higher RER at night during feeding, is necessary to proper metabolic health (65, 66). The ability to drive reduction in RER during the day at thermoneutrality speaks to the potency of DMH^TrkB^ neuronal activity in regulating fat metabolism. Acute exposure to high fat diet results in a temporary increase in locomotor activity and energy expenditure accompanied by a shift in the RER to lower values reflecting increased fat metabolism. High-fat diet results in a caloric surplus which suppresses feeding and induces increased energy expenditure. Additionally, excess lipids are targeted for use as a metabolic substrate which results in a reduced RER. Thus, DMH^TrkB→PVH/POA^ neurons may function to prime whole-animal physiology and behavior in order to rebalance a positive energy state.

The DMH regulates a plethora of vital processes (7, 10–12, 14) which is mirrored both by the complexity of the physical structure of the DMH and by its neuronal heterogeneity. Consistent with previous studies investigating the function and neurocircuitry of other populations of DMH neurons (8, 14, 20, 24, 67), we see that thermogenic DMH^TrkB→RPa^ neurons appear to be concentrated in the anterior and dorsal regions of the DMH, while food-regulatory DMH^TrkB→PVH/POA^ neurons are localized in the middle and ventral DMH. A key question remains, what are the identities of DMH^TrkB^ neurons that project either to the RPa or the PVH/POA? DMH neurons that express the leptin receptor (LepR) (21, 29), prolactin-releasing peptide (PrRP) (67), or bombesin-like receptor 3 (Brs3) (20) have been shown to promote thermogenesis, however their additional effects on heart rate and blood pressure make it unlikely that DMH^TrkB→RPa^ neurons represents one of these known populations. Furthermore, we have previously established that few DMH^TrkB^ neurons respond to leptin and they do not form synaptic connections with AgRP-expressing neurons in the arcuate (26) as have been reported for DMH^LepR^ neurons that acutely regulate feeding (24). Finally, some DMH^TrkB^ neurons co-express cholecystokinin (CCK), and loss of TrkB in DMH^CCK^ neurons results in weight gain attributed to hypercortisolism without altering food intake, energy expenditure, or locomotor activity (68). Therefore, DMH^TrkB/CCK^ neurons are unlikely to represent the DMH^TrkB^ populations that we find project to the RPa or the PVH/POA and the molecular identity of DMH^TrkB^ neuron populations remains unclear. Future experiments that take advantage of recent advances in single cell sequencing technology will undoubtably provide valuable information regarding markers to distinguish different DMH^TrkB^ populations.

In summary, our data indicate that DMH^TrkB^ neurons act through multiple neurocircuits to regulate disparate processes that are key to maintaining energy homeostasis. DMH^TrkB^ neurons suppress feeding, promote adaptive thermogenesis, enhance physical activity, and reduce the respiratory quotient. Notably, DMH^TrkB^ neurons can induce increased energy expenditure and fat metabolism without affecting cardiovascular physiology. Additionally, we reveal that separate populations of DMH^TrkB^ neurons are responsible for modulating thermogenesis or metabolism and feeding through projections to the RPa or the POA and PVH respectively. Our results highlight the importance of DMH^TrkB^ neuronal activity in energy balance and provide insight to the neural mechanisms of their influence which may be expected to inform efforts to target these neurons for the development anti-obesity therapies.

## Methods

### Mice

Mice were maintained on a C57BL6/J background in a climate-controlled facility at 22°C on a 12h /12h light-dark cycle with *ad libitum* access to water and chow diet (Teklad Rodent Diet 2920) unless otherwise specified. The *Ai9/+* mouse strain [Gt(ROSA)26Sortm9(CAG-tdTomato)Hze/J; stock no. 007909] was acquired from Jackson Laboratory. The *Ntrk2*^*Cre-ER/+*^ (also known as TrkB^CreER^) mouse strain was generously provided by Dr. David Ginty (Harvard Medical School). Male and female mice aged 6-12 weeks old were used for initial investigations of chemogenetic activation or inhibition of global DMH^TrkB^ neurons. Both males and females showed similar phenotypes, thus only female mice were used for subsequent projection specific manipulation of DMH^TrkB^ neuronal activity. Nuclear localization of Cre recombinase in *Ntrk2*^*Cre-ER/+*^ mice was induced by tamoxifen (20 mg/ml in corn oil, Sigma-Aldrich T-5648) administration for 6 days [3 mg per animal per day, intraperitoneally (i.p.)] except where otherwise indicated. For DREADD experiments, mice were injected i.p. with CNO (1.5 mg/kg, Cayman Chemical 16882).

### Viruses and Plasmid construction

AAV2-CMV-GFP (4.4×10^12^ vg/ml, UNC Vector Core), AAV2-CAG-FLEX-tdTomato (4.8×10^12^ vg/ml, UNC Vector Core), AAV8-hSyn-DIO-hM3-mCherry (4×10^12^ vg/ml, Addgene 44362), AAV8-hSyn-hM4-mCherry (6×10^12^ vg/ml, UNC Vector Core), AAV8-hSyn-mCherry (2.1×10^13^ vg/mL, Addgene). AAV2-retro-EF1a-DIO-FLPo (2.6×10^13^ vg/ml) was packaged by Vigene Biosciences from pAAV-EF1a-DIO-FLPo-WPRE-hGHpA (gift from Li Zhang, Addgene 87306). AAV2-fDIO-mCherry was previously described(42) (4.2×10^12^ vg/ml, packaged by Vigene), and AAV2-fDIO-hM3-mCherry (6.64×10^12^ vg/ml, packaged by Vigene) was similarly constructed: pAAV-hSyn-DIO-hM3-mCherry (gift from Brian Roth, Addgene 44361) and pAAV-EF1a-fDIO-EYFP (gift from Karl Deisseroth, Addgene 55461) were digested with AscI and NheI, and the hM3-mCherry fragment was ligated into pAAV-EF1a-fDIO backbone. The resulting pAAV-EF1a-fDIO-hM3-mCherry plasmid was confirmed by sequencing and additional digestion with XmaI prior to commercial packaging. Herpes simplex virus (HSV)129ΔTK-tdTomato as obtained from the Center for Neuroanatomy with Neurotropic Viruses.

### Immunohistochemistry

For Fos quantification in *Ntrk2*^*Cre-ER/+*^;*Ai9/+* animals, labeling of TrkB+ neurons was induced with tamoxifen as described above. One week after the last tamoxifen injection, mice were individually placed in fresh cages in either warm (39°C), cold (10°C), or thermoneutral (30°C) ambient temperatures 2 hours prior to transcardial perfusion with 4% paraformaldehyde (PFA). Brains were dissected, post-fixed in 4%PFA overnight at 4°C, and cryoprotected in 30% sucrose for 3 days prior to collection of 40 µm floating coronal sections. For Fos staining to confirm projection-specific activation of DMH^TrkB→RPa^ neurons, mice expressing either mCherry or hM3 in DMH^TrkB→RPa^ neurons were treated with CNO 2 hours prior to dissection and fixation of the brain. Sections were permeabilized in 1% Triton-×100 (T-×100), incubated in blocking buffer (10% horse serum, 1% bovine serum albumin (BSA), 0.1% T-×100 in PBS) for one hour at room temperature (RT) followed by incubation with primary antibody, rabbit anti-cFos (1:10,000 Synaptic System #226003) in 1% horse serum, 0.1%BSA, 0.1%T-×100 in PBS overnight at 4°C. The next day, sections were washed in PBS before incubation with secondary antibody (1:800, 488-donkey anti-rabbit, Jackson ImmunoResearch, 171-545-152) in blocking buffer for 1 hour at RT. Sections were washed again in PBS before staining for 20 minutes in DAPI (5 µg/mL, Roche 10236276001) in PBS. Sections were then mounted on Superfrost plus microscope slides (Fisher, 12-550-15) and sealed with mounting media (2.5% PVA-DABCO) and glass coverslips.

For analysis of anterograde tracing in BAT and heart tissues, mice were perfused with PBS followed by 4% PFA, and brown adipose tissue and heart were dissected and fixed overnight in 4% PFA. Tissues were cryoprotected in 30% sucrose, embedded in OCT, and frozen before collecting 20 µm sections with a cryostat. Sections were permeabilized in 1% T-×100 for 9 minutes, blocked in 5% horse serum/1% bovine serum albumin (BSA)/0.1% T-×100 for 1 hour, and incubated with primary antibodies (rabbit anti-UCP1, 1:750, Abcam ab10983, mouse anti-tyrosine hydroxylase, 1:1,500, Sigma-Aldrich T1299) in blocking buffer overnight at 4°C. Sections were washed in PBS and incubated with secondary antibodies (1:800, Jackson ImmunoResearch) in blocking buffer for 1 hour at RT. Sections were washed with PBS and stained with DAPI and sealed with mounting media (2.5%PVA-DAPCO) and glass coverslips.

Images representing 1µm optical slices were taken using a Nikon C2+ confocal microscope equipped with 405, 488, 561, and 640nm coherent lasers. For Fos quantification of DMH^TrkB^ neurons, every third brain section was imaged bilaterally for a total of 7 sections spanning 520µm of the DMH. Cell counts were performed using ImageJ Cell counter (NIH). Averages of TrkB+ and TrkB+/Fos+ neurons were taken for the first two sections representing the anterior DMH, the 3^rd^-5^th^ sections representing the middle DMH, and the last two sections representing the posterior DMH.

### Stereotaxic surgery

*Ntrk2*^*Cre-ER/+*^ mice at 6-8 weeks of age were anesthetized with isoflurane and mounted on a stereotaxic frame (Kopf Model 960). An incision through the skin along the midline of the head was made to expose the skull and the location of Bregma was determined as the reference point for the injection site coordinates [anterior-posterior (AP)/ mediolateral (ML)/ dorsoventral (DV)]: POA (AP, +0.60 mm/ ML, ±0.35 mm/ DV, −5.70 mm), PVH (AP, −0.50 mm/ ML, ±0.35 mm/ DV, −5.30 mm), DMH (AP, −1.50 mm/ ML, ±0.35 mm, DV, −5.70 mm), RPa (AP, −6.00 mm/ ML, 0.00 mm/ DV, −6.35 mm). A small hole was drilled through the skull at the injection site, the dura was gently peeled away, and a 2.5-µl syringe (Hamilton 7632-01) with 33G needle was slowly lowered into the brain. Virus was infused at a rate of 25 nl/min with a micropump and controller (World Precision Instruments, UMP3 pump and Micro4 controller) for a total of 250 nl (DMH) or 150 nl (POA, PVH, and RPa). Mice were allowed to recover for 1 week before induction with tamoxifen as described above, and experiments were conducted 1 week following the last tamoxifen injection. CTB-488 and CTB-647 (Invitrogen) were stereotaxically injected into a brain site at 1 mg/ml.

For polysynaptic anterograde tracing with HSV129ΔTK-tt, *Ntrk2*^*CreER/+*^ mice were given daily injections of tamoxifen (3mg/animal) starting 2 days prior to stereotaxic viral injection of HSV129ΔTK-TT (150nl, 2.2×10^10^ pfu/mL) unilaterally into the DMH, and continuing on postoperative days 1-4.

### Core body temperature measurements

Animals were allowed to acclimate for 6 days to thermoneutral (30°C) or cold (10°C) housing for excitatory hM3 or inhibitory hM4 DREADD experiments respectively. During this time animals were also conditioned to mock i.p. injections, and temperature measurements with a digital thermometer (Fisher Traceable Type-K thermometer) with rodent rectal temperature probe (World Precision Instruments, RET-3). For experiments performed in a cold environment, animals were housed individually. On the day of the experiment, rectal temperature was recorded immediately prior to i.p. injection of vehicle. Rectal temperature was recorded every 20 minutes for 1 hour. After the collection of the last timepoint following vehicle injection, another baseline measurement was recorded immediately prior to injection with CNO, and rectal temperature was acquired every 20 minutes for 1 hour for mice housed at thermoneutrality and for 1.5 hours for mice housed at 10°C. This procedure was repeated for two additional days and the average of the measurements for three days was determined for each animal.

### iBAT temperature measurements

Animals were allowed to acclimate to thermoneutral housing and handling 6 days prior to experimentation. The day before testing, mice were anesthetized and the hair on their backs and above their shoulders was shaved. On the day of experimentation, a thermal camera (FLIR E53sc) was mounted above a cage with mice expressing either control (mCherry) or excitatory (hM3) virus in DMH^TrkB^ neurons. Thermal images were collected 1 hour after i.p. injection of vehicle or CNO and regions above the shoulders (above interscapular BAT), lumbar area (body), and tail were analyzed using Research IR software (FLIR). The average of 3 days of testing was determined for each animal.

### Physiological measurements

Locomotor activity and oxygen consumption in individual mice were measured using a comprehensive lab animal monitoring system (CLAMS, Columbus Instruments). Mice were previously conditioned for at least 1 week to thermoneutrality (for excitatory DREADD experiments) and 10°C (for inhibitory DREADD experiments) and allowed to acclimate to metabolic chambers for at least 24 hours prior to experimentation. Mice received a vehicle i.p. injection in the morning (between 8-9am) and a second i.p injection with CNO (between 12-1pm). Physical activity and metabolic data were collected for 4 hours following treatment with either vehicle or CNO. This paradigm was repeated for a second day and the data for each animal was averaged for days 1 and 2.

Heart rate and blood pressure were recorded using a tail cuff system (MC4000, Hatteras Instruments). Animals were acclimated to handling and restraint in the tail cuffs for 5 days prior to experimentation. The platform was set to 35°C, with 10 preliminary measurement cycles were followed by 15 cycles of measurement. Animals were placed in the tail cuffs 30 minutes after i.p injection with vehicle or CNO. Vehicle and CNO measurements were collected on different days at the same time of day to minimize the effect of stress due to restraint in the tail cuffs.

Nocturnal food intake studies were performed as previously described(26). Mice were acclimated to handling and single housing for 3 days prior to experimentation. Food intake was monitored for 4 nights following treatment with either vehicle (nights 1 and 2) or CNO (nights 3 and 4). On the day of the day of the experiment, food was removed from the cage 3 hours prior to lights-off, and animals were given an i.p. injection of vehicle or CNO 10 minutes before lights-off. At the onset of lights-off, two food pellets (Teklad Rodent Diet 2920X; 3.1 kcal/g energy density) were weighed (with 0.01g precision) and placed in the cage with the mouse. Food was weighed again every hour for 4 hours after lights-off. Food intake for the two nights following either vehicle or CNO injection were averaged for each animal.

### Quantitative RT-PCR

For measurement of *Ucp1* levels in BAT following activation of DMH^TrkB^ neurons, BAT tissue was dissected 2 hours after injection of CNO and snap-frozen in liquid nitrogen. RNA was extracted from 15-20 mg of BAT with TRIzol (Invitrogen) and chloroform and precipitated with isopropanol. RNA was reverse-transcribed with M-MuLV reverse transcriptase (New England Biolabs, M0253) and random hexomers (ThermoFisher N8080127), and cDNA was diluted (1:5) prior to quantitative PCR (qPCR) analysis. qPCR was carried out using SYBR green mix (Roche) in an StepOne cycler (Applied Biosystems), using the primers: UCP1-F: 5′-GATGGTGAACCCGACAACTT-3′, UCP1-R: 5′-CTGAAACTCCGGCTGAGAAG-3′, 18S rRNA-F: 5′-CGCCGCTAGAGGTGAAATTC-3′, 18S rRNA-R: 5′-TTGGCAAATGCTTTCGCTC-3′. Fold change in UCP-1 transcript levels was calculated using the −2ΔΔCt method, normalizing to 18S rRNA transcript.

### Statistical analyses

All data are presented as means ± standard error of the mean (S.E.M.). Graphs and statistical analyses were generated using GraphPad Prism software. Statistical significance was determined using either two-tailed unpaired Student’s t-test, or two-way ANOVA when comparing more than two groups. All experiments were performed at least 3 times independently or in 3 separate animals. All statistical analyses are listed in Supplemental Table 1A-D.

### Study approval

All experiments were performed in accordance with relevant guidelines and regulations regarding the use of experimental animals. The Animal Care and Use Committees at The Scripps Research Institute Florida approved all animal procedures used in this study.

## Author contributions

J.H. designed and performed the majority of the experiments, analyzed all the data, and wrote the paper. G.-Y.L. performed stereotaxic injection of AAV2-FLEX-TdTomato for anterograde tracing. B.X. provided guidance and resources and edited the paper.

## Acknowledgements

This work was supported by grants from the National Institutes of Health to BX (R01 DK105954 and R01 DK103335) and JH (F32 NS106810). HSV129ΔTK-TT was provided by the Center for Neuroanatomy with Neurotropic Viruses, which was supported by an NIH grant (P40 RR018604).

## References

1. Hruby A, and Hu FB. The Epidemiology of Obesity: A Big Picture. PharmacoEconomics. 2015;33(7):673–89.

2. Hall KD, and Kahan S. Maintenance of Lost Weight and Long-Term Management of Obesity. The Medical clinics of North America. 2018;102(1):183–97.

3. Knowler WC, Fowler SE, Hamman RF, Christophi CA, Hoffman HJ, Brenneman AT, et al. 10-year follow-up of diabetes incidence and weight loss in the Diabetes Prevention Program Outcomes Study. Lancet (London, England). 2009;374(9702):1677–86.

4. Lotfi K, Palmer K, and Apovian CM. Case Study: Weight loss in a patient with type 2 diabetes: Challenges of diabetes management. Obesity (Silver Spring, Md). 2015;23 Suppl 1:S11–2.

5. Chen KY, Brychta RJ, Abdul Sater Z, Cassimatis TM, Cero C, Fletcher LA, et al. Opportunities and challenges in the therapeutic activation of human energy expenditure and thermogenesis to manage obesity. The Journal of biological chemistry. 2020;295(7):1926–42.

6. Kalil GZ, and Haynes WG. Sympathetic nervous system in obesity-related hypertension: mechanisms and clinical implications. Hypertension research : official journal of the Japanese Society of Hypertension. 2012;35(1):4–16.

7. Lee SJ, Kirigiti M, Lindsley SR, Loche A, Madden CJ, Morrison SF, et al. Efferent projections of neuropeptide Y-expressing neurons of the dorsomedial hypothalamus in chronic hyperphagic models. The Journal of comparative neurology. 2013;521(8):1891–914.

8. Zhao ZD, Yang WZ, Gao C, Fu X, Zhang W, Zhou Q, et al. A hypothalamic circuit that controls body temperature. Proceedings of the National Academy of Sciences of the United States of America. 2017;114(8):2042–7.

9. Chen M, Shrestha YB, Podyma B, Cui Z, Naglieri B, Sun H, et al. Gsα deficiency in the dorsomedial hypothalamus underlies obesity associated with Gsα mutations. The Journal of clinical investigation. 2017;127(2):500–10.

10. Aston-Jones G, Chen S, Zhu Y, and Oshinsky ML. A neural circuit for circadian regulation of arousal. Nature neuroscience. 2001;4(7):732–8.

11. Chen KS, Xu M, Zhang Z, Chang WC, Gaj T, Schaffer DV, et al. A Hypothalamic Switch for REM and Non-REM Sleep. Neuron. 2018;97(5):1168–76.e4.

12. Kataoka N, Hioki H, Kaneko T, and Nakamura K. Psychological stress activates a dorsomedial hypothalamus-medullary raphe circuit driving brown adipose tissue thermogenesis and hyperthermia. Cell metabolism. 2014;20(2):346–58.

13. Marchant NJ, Furlong TM, and McNally GP. Medial dorsal hypothalamus mediates the inhibition of reward seeking after extinction. The Journal of neuroscience : the official journal of the Society for Neuroscience. 2010;30(42):14102–15.

14. Nakamura Y, Nakamura K, Matsumura K, Kobayashi S, Kaneko T, and Morrison SF. Direct pyrogenic input from prostaglandin EP3 receptor-expressing preoptic neurons to the dorsomedial hypothalamus. The European journal of neuroscience. 2005;22(12):3137–46.

15. Tan CL, Cooke EK, Leib DE, Lin YC, Daly GE, Zimmerman CA, et al. Warm-Sensitive Neurons that Control Body Temperature. Cell. 2016;167(1):47–59.e15.

16. Nakamura K, Matsumura K, Hubschle T, Nakamura Y, Hioki H, Fujiyama F, et al. Identification of sympathetic premotor neurons in medullary raphe regions mediating fever and other thermoregulatory functions. The Journal of neuroscience : the official journal of the Society for Neuroscience. 2004;24(23):5370–80.

17. Schneeberger M, Parolari L, Das Banerjee T, Bhave V, Wang P, Patel B, et al. Regulation of Energy Expenditure by Brainstem GABA Neurons. Cell. 2019;178(3):672–85.e12.

18. Cao WH, and Morrison SF. Glutamate receptors in the raphe pallidus mediate brown adipose tissue thermogenesis evoked by activation of dorsomedial hypothalamic neurons. Neuropharmacology. 2006;51(3):426–37.

19. Zaretskaia MV, Zaretsky DV, Shekhar A, and DiMicco JA. Chemical stimulation of the dorsomedial hypothalamus evokes non-shivering thermogenesis in anesthetized rats. Brain research. 2002;928(1-2):113–25.

20. Pinol RA, Zahler SH, Li C, Saha A, Tan BK, Skop V, et al. Brs3 neurons in the mouse dorsomedial hypothalamus regulate body temperature, energy expenditure, and heart rate, but not food intake. Nature neuroscience. 2018;21(11):1530–40.

21. Simonds SE, Pryor JT, Ravussin E, Greenway FL, Dileone R, Allen AM, et al. Leptin mediates the increase in blood pressure associated with obesity. Cell. 2014;159(6):1404–16.

22. Bellinger LL, and Bernardis LL. The dorsomedial hypothalamic nucleus and its role in ingestive behavior and body weight regulation: lessons learned from lesioning studies. Physiology & behavior. 2002;76(3):431–42.

23. Otgon-Uul Z, Suyama S, Onodera H, and Yada T. Optogenetic activation of leptin- and glucose-regulated GABAergic neurons in dorsomedial hypothalamus promotes food intake via inhibitory synaptic transmission to paraventricular nucleus of hypothalamus. Molecular metabolism. 2016;5(8):709–15.

24. Garfield AS, Shah BP, Burgess CR, Li MM, Li C, Steger JS, et al. Dynamic GABAergic afferent modulation of AgRP neurons. Nature neuroscience. 2016;19(12):1628–35.

25. Rau AR, and Hentges ST. GABAergic Inputs to POMC Neurons Originating from the Dorsomedial Hypothalamus Are Regulated by Energy State. The Journal of neuroscience : the official journal of the Society for Neuroscience. 2019;39(33):6449–59.

26. Liao GY, Kinney CE, An JJ, and Xu B. TrkB-expressing neurons in the dorsomedial hypothalamus are necessary and sufficient to suppress homeostatic feeding. Proceedings of the National Academy of Sciences of the United States of America. 2019;116(8):3256–61.

27. Chen J, Scott KA, Zhao Z, Moran TH, and Bi S. Characterization of the feeding inhibition and neural activation produced by dorsomedial hypothalamic cholecystokinin administration. Neuroscience. 2008;152(1):178–88.

28. Sonoyama T, Stadler LKJ, Zhu M, Keogh JM, Henning E, Hisama F, et al. Human BDNF/TrkB variants impair hippocampal synaptogenesis and associate with neurobehavioural abnormalities. Scientific reports. 2020;10(1):9028.

29. Zhang Y, Kerman IA, Laque A, Nguyen P, Faouzi M, Louis GW, et al. Leptin-receptor-expressing neurons in the dorsomedial hypothalamus and median preoptic area regulate sympathetic brown adipose tissue circuits. The Journal of neuroscience : the official journal of the Society for Neuroscience. 2011;31(5):1873–84.

30. Rutlin M, Ho CY, Abraira VE, Cassidy C, Bai L, Woodbury CJ, et al. The cellular and molecular basis of direction selectivity of Adelta-LTMRs. Cell. 2014;159(7):1640–51.

31. Madisen L, Zwingman TA, Sunkin SM, Oh SW, Zariwala HA, Gu H, et al. A robust and high-throughput Cre reporting and characterization system for the whole mouse brain. Nature neuroscience. 2010;13(1):133–40.

32. Greenberg ME, Ziff EB, and Greene LA. Stimulation of neuronal acetylcholine receptors induces rapid gene transcription. Science (New York, NY). 1986;234(4772):80–3.

33. Alexander GM, Rogan SC, Abbas AI, Armbruster BN, Pei Y, Allen JA, et al. Remote control of neuronal activity in transgenic mice expressing evolved G protein-coupled receptors. Neuron. 2009;63(1):27–39.

34. Fischer AW, Cannon B, and Nedergaard J. Optimal housing temperatures for mice to mimic the thermal environment of humans: An experimental study. Molecular metabolism. 2017.

35. Meyer CW, Ootsuka Y, and Romanovsky AA. Body Temperature Measurements for Metabolic Phenotyping in Mice. Frontiers in physiology. 2017;8:520.

36. Takahashi TM, Sunagawa GA, Soya S, Abe M, Sakurai K, Ishikawa K, et al. A discrete neuronal circuit induces a hibernation-like state in rodents. Nature. 2020.

37. Brito NA, Brito MN, and Bartness TJ. Differential sympathetic drive to adipose tissues after food deprivation, cold exposure or glucoprivation. American journal of physiology Regulatory, integrative and comparative physiology. 2008;294(5):R1445–52.

38. Tupone D, Madden CJ, and Morrison SF. Autonomic regulation of brown adipose tissue thermogenesis in health and disease: potential clinical applications for altering BAT thermogenesis. Frontiers in neuroscience. 2014;8:14.

39. An JJ, Liao GY, Kinney CE, Sahibzada N, and Xu B. Discrete BDNF Neurons in the Paraventricular Hypothalamus Control Feeding and Energy Expenditure. Cell metabolism. 2015;22(1):175–88.

40. Yeo GS, Connie Hung CC, Rochford J, Keogh J, Gray J, Sivaramakrishnan S, et al. A de novo mutation affecting human TrkB associated with severe obesity and developmental delay. Nature neuroscience. 2004;7(11):1187–9.

41. Xu B, Goulding EH, Zang K, Cepoi D, Cone RD, Jones KR, et al. Brain-derived neurotrophic factor regulates energy balance downstream of melanocortin-4 receptor. Nature neuroscience. 2003;6(7):736–42.

42. An JJ, Kinney CE, Tan JW, Liao GY, Kremer EJ, and Xu B. TrkB-expressing paraventricular hypothalamic neurons suppress appetite through multiple neurocircuits. Nature communications. 2020;11(1):1729.

43. Zhang N, Yang L, Guo L, and Bi S. Activation of Dorsomedial Hypothalamic Neurons Promotes Physical Activity and Decreases Food Intake and Body Weight in Zucker Fatty Rats. Frontiers in molecular neuroscience. 2018;11:179.

44. Yu S, Qualls-Creekmore E, Rezai-Zadeh K, Jiang Y, Berthoud HR, Morrison CD, et al. Glutamatergic Preoptic Area Neurons That Express Leptin Receptors Drive Temperature-Dependent Body Weight Homeostasis. The Journal of neuroscience : the official journal of the Society for Neuroscience. 2016;36(18):5034–46.

45. Kong D, Tong Q, Ye C, Koda S, Fuller PM, Krashes MJ, et al. GABAergic RIP-Cre neurons in the arcuate nucleus selectively regulate energy expenditure. Cell. 2012;151(3):645–57.

46. Madden CJ, and Morrison SF. Neurons in the paraventricular nucleus of the hypothalamus inhibit sympathetic outflow to brown adipose tissue. American journal of physiology Regulatory, integrative and comparative physiology. 2009;296(3):R831–43.

47. Miao Y, Wu W, Dai Y, Maneix L, Huang B, Warner M, et al. Liver X receptor β controls thyroid hormone feedback in the brain and regulates browning of subcutaneous white adipose tissue. Proceedings of the National Academy of Sciences of the United States of America. 2015;112(45):14006–11.

48. Bovetto S, Rouillard C, and Richard D. Role of CRH in the effects of 5-HT-receptor agonists on food intake and metabolic rate. The American journal of physiology. 1996;271(5 Pt 2):R1231–8.

49. Villarroya F, and Vidal-Puig A. Beyond the sympathetic tone: the new brown fat activators. Cell metabolism. 2013;17(5):638–43.

50. Lebow MA, and Chen A. Overshadowed by the amygdala: the bed nucleus of the stria terminalis emerges as key to psychiatric disorders. Molecular psychiatry. 2016;21(4):450–63.

51. Kodani S, Soya S, and Sakurai T. Excitation of GABAergic Neurons in the Bed Nucleus of the Stria Terminalis Triggers Immediate Transition from Non-Rapid Eye Movement Sleep to Wakefulness in Mice. The Journal of neuroscience : the official journal of the Society for Neuroscience. 2017;37(30):7164–76.

52. Hua R, Wang X, Chen X, Wang X, Huang P, Li P, et al. Calretinin Neurons in the Midline Thalamus Modulate Starvation-Induced Arousal. Current biology : CB. 2018;28(24):3948–59.e4.

53. Li MM, Madara JC, Steger JS, Krashes MJ, Balthasar N, Campbell JN, et al. The Paraventricular Hypothalamus Regulates Satiety and Prevents Obesity via Two Genetically Distinct Circuits. Neuron. 2019;102(3):653–67.e6.

54. Xu Y, Wu Z, Sun H, Zhu Y, Kim ER, Lowell BB, et al. Glutamate mediates the function of melanocortin receptor 4 on Sim1 neurons in body weight regulation. Cell metabolism. 2013;18(6):860–70.

55. Huang EJ, and Reichardt LF. Neurotrophins: roles in neuronal development and function. Annual review of neuroscience. 2001;24:677–736.

56. Lin PY, Kavalali ET, and Monteggia LM. Genetic Dissection of Presynaptic and Postsynaptic BDNF-TrkB Signaling in Synaptic Efficacy of CA3-CA1 Synapses. Cell reports. 2018;24(6):1550–61.

57. Zakharenko SS, Patterson SL, Dragatsis I, Zeitlin SO, Siegelbaum SA, Kandel ER, et al. Presynaptic BDNF required for a presynaptic but not postsynaptic component of LTP at hippocampal CA1-CA3 synapses. Neuron. 2003;39(6):975–90.

58. Bayazitov IT, Richardson RJ, Fricke RG, and Zakharenko SS. Slow presynaptic and fast postsynaptic components of compound long-term potentiation. The Journal of neuroscience : the official journal of the Society for Neuroscience. 2007;27(43):11510–21.

59. Kroeger D, Absi G, Gagliardi C, Bandaru SS, Madara JC, Ferrari LL, et al. Galanin neurons in the ventrolateral preoptic area promote sleep and heat loss in mice. Nature communications. 2018;9(1):4129.

60. Abbott SB, Machado NL, Geerling JC, and Saper CB. Reciprocal Control of Drinking Behavior by Median Preoptic Neurons in Mice. The Journal of neuroscience : the official journal of the Society for Neuroscience. 2016;36(31):8228–37.

61. Wei YC, Wang SR, Jiao ZL, Zhang W, Lin JK, Li XY, et al. Medial preoptic area in mice is capable of mediating sexually dimorphic behaviors regardless of gender. Nature communications. 2018;9(1):279.

62. Yu S, François M, Huesing C, and Münzberg H. The Hypothalamic Preoptic Area and Body Weight Control. Neuroendocrinology. 2018;106(2):187–94.

63. Moffitt JR, Bambah-Mukku D, Eichhorn SW, Vaughn E, Shekhar K, Perez JD, et al. Molecular, spatial, and functional single-cell profiling of the hypothalamic preoptic region. Science (New York, NY). 2018;362(6416).

64. Hrvatin S, Sun S, Wilcox OF, Yao H, Lavin-Peter AJ, Cicconet M, et al. Neurons that regulate mouse torpor. Nature. 2020.

65. Chaix A, Lin T, Le HD, Chang MW, and Panda S. Time-Restricted Feeding Prevents Obesity and Metabolic Syndrome in Mice Lacking a Circadian Clock. Cell metabolism. 2019;29(2):303–19.e4.

66. Müller MJ, Enderle J, Pourhassan M, Braun W, Eggeling B, Lagerpusch M, et al. Metabolic adaptation to caloric restriction and subsequent refeeding: the Minnesota Starvation Experiment revisited. The American journal of clinical nutrition. 2015;102(4):807–19.

67. Dodd GT, Worth AA, Nunn N, Korpal AK, Bechtold DA, Allison MB, et al. The thermogenic effect of leptin is dependent on a distinct population of prolactin-releasing peptide neurons in the dorsomedial hypothalamus. Cell metabolism. 2014;20(4):639–49.

68. Geibel M, Badurek S, Horn JM, Vatanashevanopakorn C, Koudelka J, Wunderlich CM, et al. Ablation of TrkB signalling in CCK neurons results in hypercortisolism and obesity. Nature communications. 2014;5:3427.

