## Supplemental Data for "Discrete TrkB-expressing neurons of the dorsomedial hypothalamus regulate feeding and thermogenesis"

**Supplemental Information**

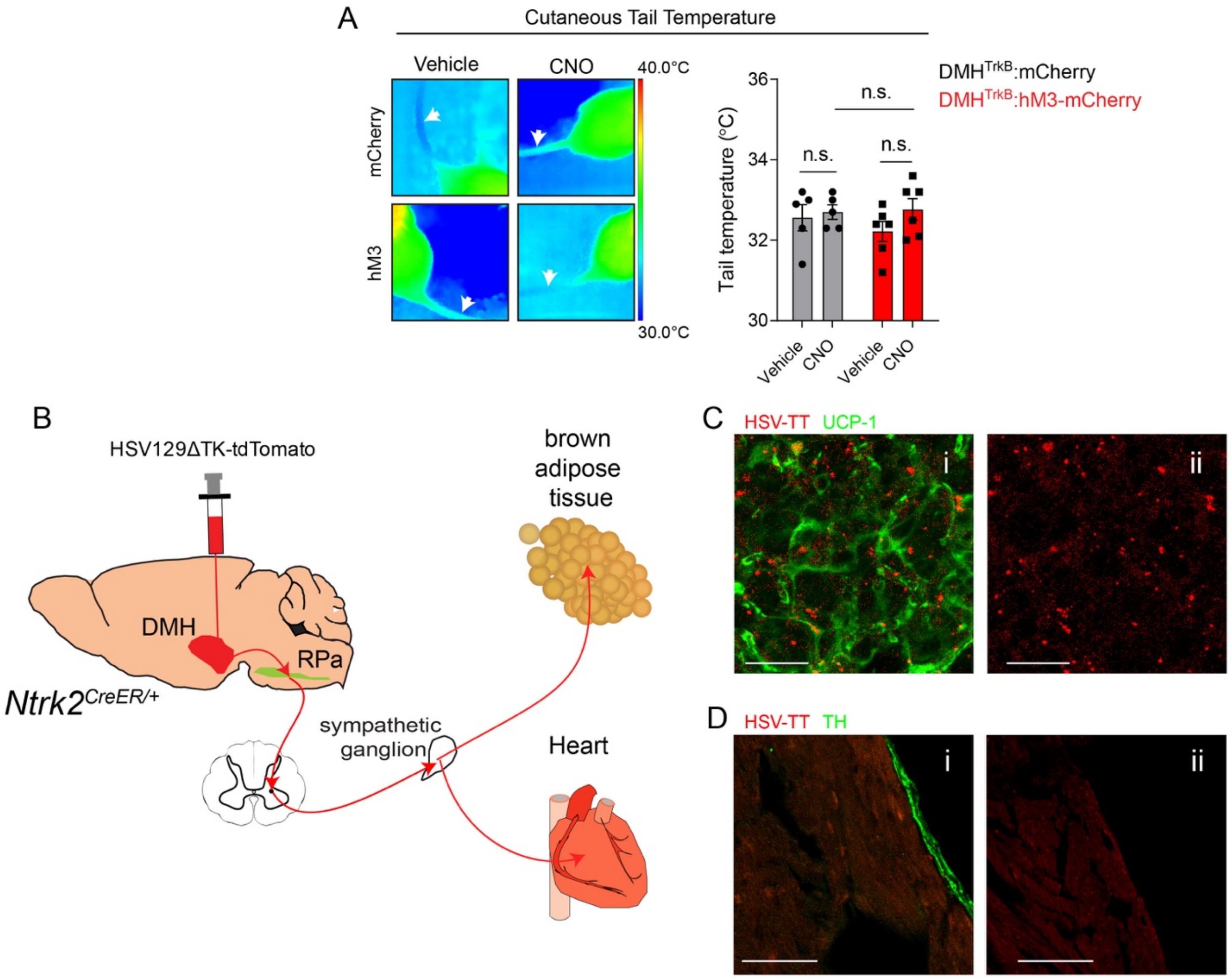

**Supplemental Figure 1. DMH$^{TrkB}$ neurons do not influence vasoconstriction and form polysynaptic connections with BAT but not heart.** (**A**) Activation of DMH$^{TrkB}$ neurons did not alter cutaneous tail temperature (indicator of vasoconstriction or vasodilation) as measured by thermal imaging (*n* = 5 mCherry, 6 hM3). Two-way RM ANOVA: mCherry vs. hM3 $F_{(1,\ 18)}$ = 0.2741, p=0.6070. (**B**) Diagram of polysynaptic anterograde tracing of DMH$^{TrkB}$ neurons in *Ntrk2*$^{Cre\text{-}ER/+}$ mice. Following induction with tamoxifen, stereotactic delivery of Cre-dependent HSV129ΔTK virus expressing tdTomato into the DMH of *Ntrk2*$^{Cre\text{-}ER/+}$ mice allows for labeling of multiple orders of DMH$^{TrkB}$ neuron targets 4-5 days post-injection. (**C**) Analysis of interscapular brown adipose tissue, (**Ci**) labeled by UCP-1, showed significant (**Cii**) deposits of tdTomato indicating robust anterograde labeling from DMH$^{TrkB}$ neurons that drive thermogenesis. (**D**) Analysis of the heart revealed (**Di**) labeling of sympathetic fibers by tyrosine hydroxylase (TH), but no detectable (**Dii**) tdTomato labeling (HSV-TT) from DMH$^{TrkB}$ neurons. Values represent mean ± s.e.m., Sidak post-test (n.s. = not significant). (**C, D**) Representative images are from *n* = 4 animals. Scale bars = 20 μm (**Ci-ii**), 50 μm (**Di-ii**).

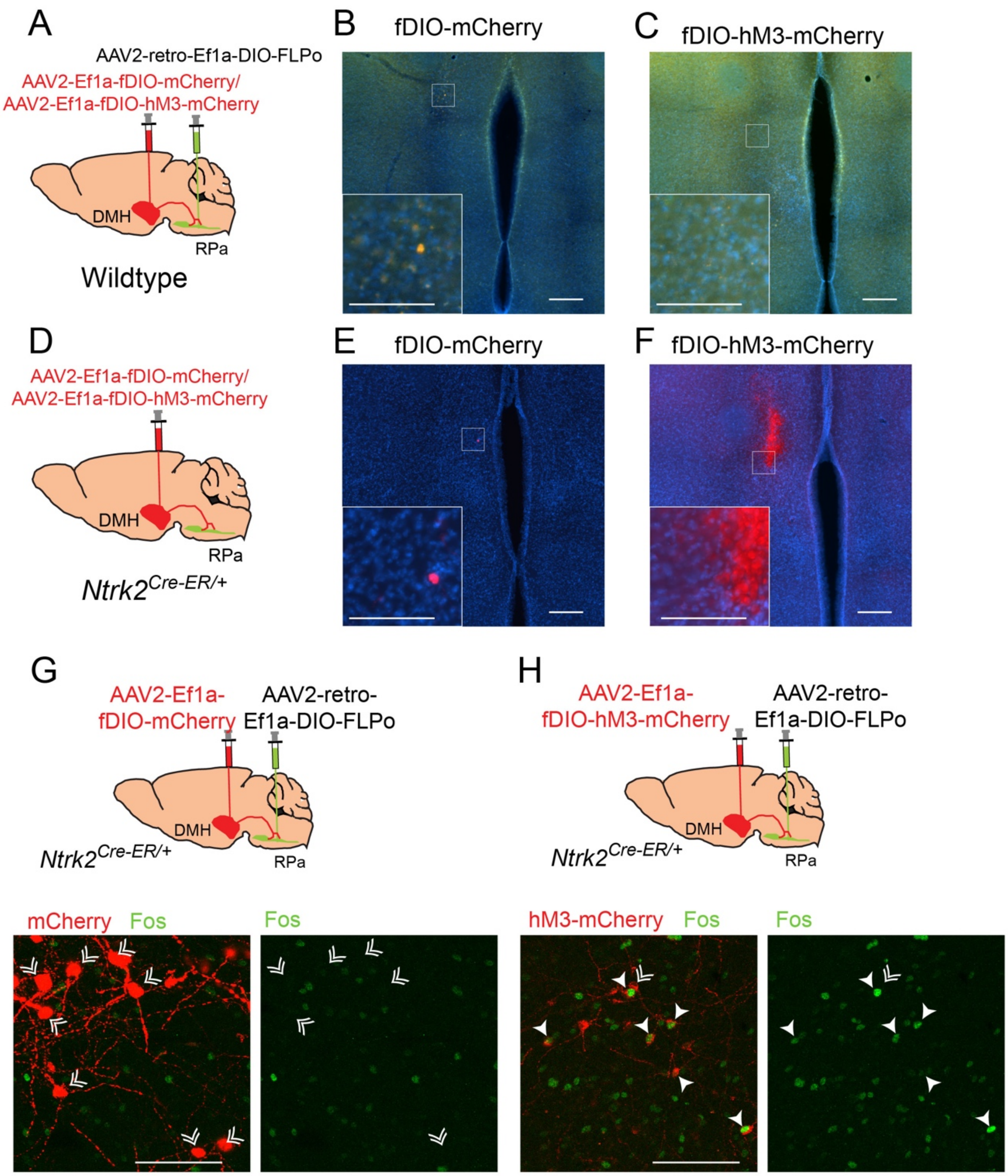

**Supplemental Figure 2. Validation of viral strategy for projection-specific expression and chemogentic excitation of DMH$^{TrkB}$ neurons.** (**A**) Schematic of stereotactic delivery of retrograde AAV expressing Cre-dependent FLPo (AAV2-retro-Ef1a-DIO-FLPo) and AAV2-CMV-GFP to the RPa, and FLP-dependent (fDIO) mCherry or hM3-mCherry expressing virus to the DMH in wildtype mice. (**B**) No expression of mCherry or (**C**) hM3-mCherry was detectable within the DMH of wildtype mice. (**D**) Injection of *Ntrk2$^{Cre-ER/+}$* with AAV expressing FLP-depended (**E**) mCherry or (**F**) hM3-mCherry does not result in expression without the additional injection of retrograde AAV-expressing Cre-dependent FLP into a DMH$^{TrkB}$ projection target. Scale bars = 200 μm, inset scale bars = 100 μm. (**G**) Expression of Fos after treatment with CNO in mice expressing mCherry or (**H**) hM3-mCherry in DMH$^{TrkB}$ neurons. Double arrowheads indicate Fos$^{-}$ DMH$^{TrkB}$ neurons. Solid arrowheads indicate Fos$^{+}$ DMH$^{TrkB}$ neurons. Scale bars = 100 μm.

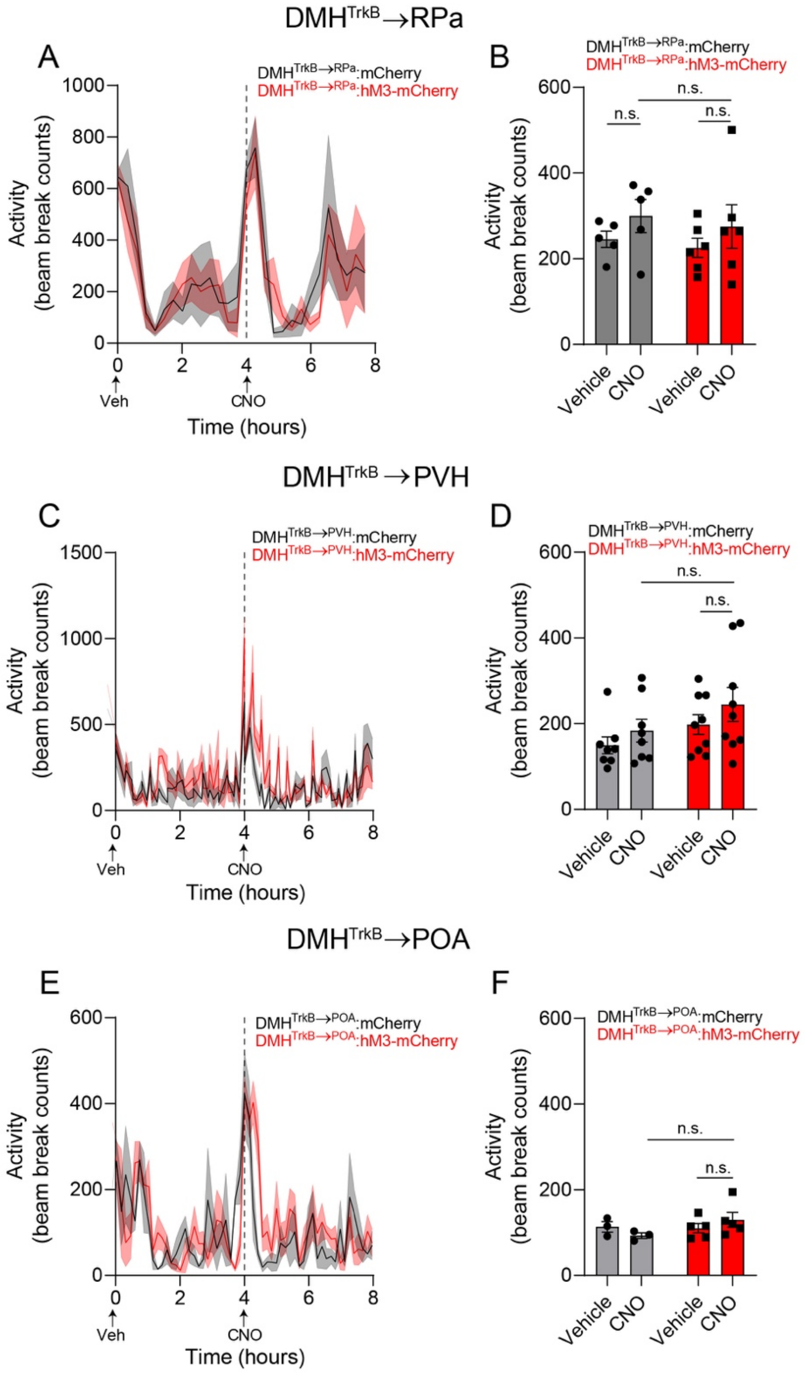

**Supplemental Figure 3. PVH-, POA-, and RPa-projecting DMH$^{TrkB}$ neurons do not regulate locomotor activity.** Mice housed at thermoneutrality were treated with vehicle (Veh) and then CNO during the light cycle. Activation of (**A**, **B**) DMH$^{TrkB\rightarrow RPa}$ neurons (*n* = 5 mCherry, 6 hM3), (**C**, **D**) DMH$^{TrkB\rightarrow PVH}$ neurons (*n* = 8 mCherry, 9 hM3), or (**E**, **F**) DMH$^{TrkB\rightarrow POA}$ neurons (*n* = 3 mCherry, 5 hM3) did not alter physical activity. (**A**, **C**, **E**) Locomotor activity after vehicle administration (hour 0-4) followed by CNO (hour 4-8). Two-way RM ANOVA: mCherry vs. hM3 (**A**) $F_{(1, 9)}$ = 0.06638, p=0.8025, (**C**) Mixed effects model: mCherry vs hM3 $F_{(1, 15)}$ = 1.478, p=0.2429, (**E**) Two-way RM ANOVA: $F_{(1, 6)}$ = 2.533, p=0.1626. (**B**, **D**, **F**) Average locomotor activity for the duration of 4 hours following either vehicle or CNO treatment. Two way RM ANOVA: mCherry vs. hM3 (**B**) $F_{(1, 9)}$ = 0.4086, p=0.5386, (**D**) $F_{(1, 15)}$ = 2.111, p=0.1669, (**F**) $F_{(1, 6)}$ = 3.189, p=0.1244. Values represent mean ± s.e.m.**,** with Sidak post-test (n.s. = not significant)

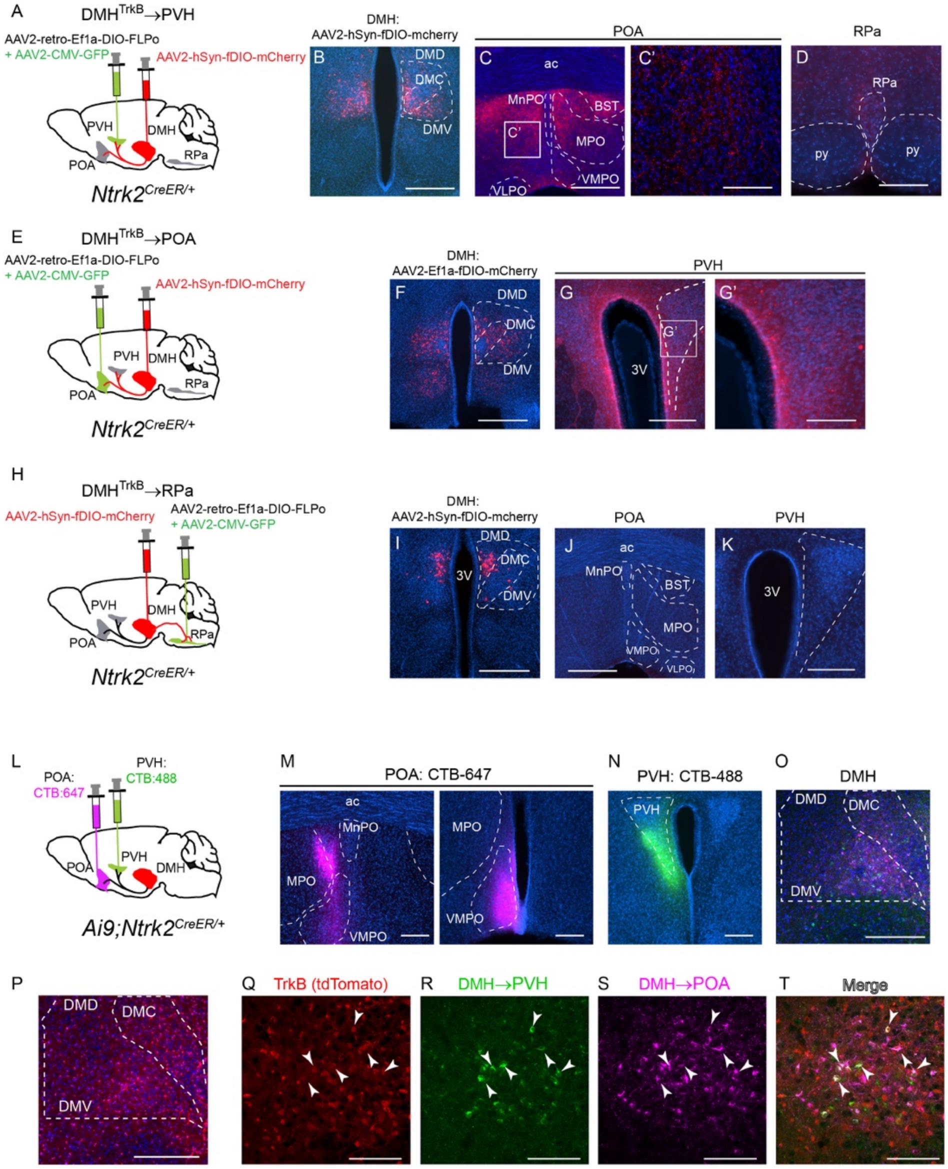

**Supplemental Figure 4: DMH$^{TrkB→PVH}$ neurons send collaterals to the POA.** (**A**-**D**) Selective viral expression of mCherry in PVH-projecting DMH$^{TrkB}$ neurons (**A**, **B**) reveals collaterals of these neurons in the POA (**C**, **C'**), but not in the RPa (**D**). (**E**-**G**) Selective viral expression of mCherry in POA-projecting DMH$^{TrkB}$ neurons (**E**, **F**) reveals collaterals of these neurons in the PVH (**G**, **G'**). (**H**-**K**) Selective viral expression of mCherry in DMH$^{TrkB→RPa}$ neurons shows that these neurons do not innervate the POA (**J**) or the PVH (**K**). (**L**) Diagram of simultaneous retrograde tracing from both the POA and the PVH in *Ai9;Ntrk2$^{Cre-ER/+}$* reporter mice. (**M**) Injection of cholera toxin subunit B Alexa Fluor 647 conjugate (CTB-647) into the POA. (**N**) Injection of CTB-488 into the PVH. **(O)** Retrograde labeling of neurons in the DMH that project to the POA (magenta) or the PVH (green). (**P**) Cre-dependent labeling of TrkB-expressing neurons in the DMH with tdTomato (red). (**Q**-**T**) Some neurons in the DMH express TrkB (red) are labeled by both CTB-488 injected into the PVH (green) and CTB-647 injected into the POA (magenta), as indicated by arrowheads. Scale bars = 500 μm in (**B**, **C**, **F**, **I**, **J**), 200 μm in (**D**, **G**, **K**, **M**-**P**), and 100 μm in (**C'**, **G'**, **Q**-**T**). Abbreviations: ac, anterior commissure; BST, bed nucleus stria terminalis; DMD, dorsal part of dorsomedial hypothalamus; DMC, central part of dorsomedial hypothalamus; DMV, ventral part of dorsomedial hypothalamus; MPO, medial preoptic area; MnPO, median preoptic area; PVH, paraventricular hypothalamus; VLPO, ventrolateral preotic area; VMPO, ventromedial preoptic area; 3V, third ventricle.

### Supplementary Table 1A: Summary of Statistical Analyses

| Figure | Sample size | Statistical test | P value |
|---|---|---|---|
| 1D | 4 mice per condition | two-way ANOVA<br>Factor 1: anterior-posterior position<br>Factor 2: temperature<br>Dunnett's multiple comparison test | position: F (1.880, 16.92) = 7.181, P = 0.2225<br>temperature F (2, 9) = 18.68, P = 0.0062<br>interaction F (4, 18) = 1.580, P= 0.0006 |
| 2B | | RM two-way ANOVA<br>Factor 1: time (post CNO)<br>Factor 2: viral expression (mcherry, hM3)<br>Sidak's multiple comparison test | time: F (1.402, 22.43) = 7.879, P = 0.0056<br>viral expression: F (1, 16) = 10.11, P = 0.0058<br>interaction: F (3, 48) = 34.69, P < 0.0001 |
| 2C | | RM two-way ANOVA<br>Factor 1: treatment (vehicle, CNO)<br>Factor 2: viral expression (mcherry, hM3)<br>Sidak's multiple comparison test | treatment: F (1, 16) = 133.3, P<0.0001<br>viral expression: F (1, 16) = 36.55, P< 0.0001<br>interaction: F (1, 16) = 151.7, P <0.0001 |
| 2D | 9 mCherry, 9 hM3 | Mixed-effects model (REML)<br>Factor 1: time (post CNO)<br>Factor 2: viral expression (mcherry, hM3) | time: F (41, 278) = 4.462, P<0.0001<br>viral expression: F (1, 16) = 15.29, P = 0.0012<br>interaction: F (41, 278) = 4.243, P < 0.0001 |
| 2E | 9 mCherry, 9 hM3 | RM two-way ANOVA<br>Factor 1: treatment (vehicle, CNO)<br>Factor 2: viral expression (mcherry, hM3)<br>Sidak's multiple comparison test | treatment: F (1, 16) = 43.35, P<0.0001<br>viral expression: F (1, 16) = 3.933, P = 0.0648<br>interaction: F (1, 16) = 29.21, P <0.0001 |
| 2F | 9 mCherry, 9 hM3 | Mixed-effects model (REML)<br>Factor 1: time (post CNO)<br>Factor 2: viral expression (mcherry, hM3) | time: F (41, 278) = 1.467, P = 0.0398<br>viral expression: F (1, 16) = 16.65, P = 0.0009<br>interaction: F (41, 278) = 2.484, P < 0.0001 |
| 2G | 9 mCherry, 9 hM3 | RM two-way ANOVA<br>Factor 1: treatment (vehicle, CNO)<br>Factor 2: viral expression (mcherry, hM3)<br>Sidak's multiple comparison test | treatment: F (1, 16) = 0.002426, P=0.0003<br>viral expression: F (1, 16) = 5.072, P = 0.0387<br>interaction: F (1, 16) = 20.88, P <0.0001 |
| 2H | 9 mCherry, 9 hM3 | Mixed-effects model (REML)<br>Factor 1: time (post CNO)<br>Factor 2: viral expression (mcherry, hM3) | time: F (41, 278) =4.667, P<0.0001<br>viral expression: F (1, 16) = 39.78, P<0.0001<br>interaction: F (41, 278) = 4.134, P < 0.0001 |
| 2I | 9 mCherry, 9 hM3 | RM two-way ANOVA<br>Factor 1: treatment (vehicle, CNO)<br>Factor 2: viral expression (mcherry, hM3)<br>Sidak's multiple comparison test | treatment: F (1, 16) = 57.24, P<0.0001<br>viral expression: F (1, 16) = 30.86, P<0.0001<br>interaction: F (1, 16) = 55.79, P <0.0001 |
| 2J | 8 mCherry, 12 hM3 | RM two-way ANOVA<br>Factor 1: treatment (vehicle, CNO)<br>Factor 2: viral expression (mcherry, hM3)<br>Sidak's multiple comparison test | treatment: F (1, 18) = 0.04648, P = 0.8317<br>viral expression: F (1, 18) = 0.6447, P = 0.4325<br>interaction: F (1, 18) = 6.618, P = 0.0192 |
| 2K | 8 mCherry, 12 hM3 | RM two-way ANOVA<br>Factor 1: treatment (vehicle, CNO)<br>Factor 2: viral expression (mcherry, hM3)<br>Sidak's multiple comparison test | treatment: F (1, 18) = 0.4427, P = 0.5143<br>viral expression: F (1, 18) = 0.01210, P = 0.9136<br>interaction: F (1, 18) = 1.586, P = 0.2240 |
| 3B | 6 mCherry, 7 hM4 | RM two-way ANOVA<br>Factor 1: time (post CNO)<br>Factor 2: viral expression (mcherry, hM4)<br>Sidak's multiple comparison test | time: F (2.993, 32.92) = 78.61, P<0.0001<br>viral expression: F (1, 11) = 20.40, P = 0.0009<br>interaction: F (4, 44) = 8.696, P < 0.0001 |

### Supplementary Table 1B: Summary of Statistical Analyses Cont.

| | | | |
|---|---|---|---|
| 3C | 6 mCherry, 7 hM4 | RM two-way ANOVA<br>Factor 1: treatment (vehicle, CNO)<br>Factor 2: viral expression (mcherry, hM4)<br>Sidak's multiple comparison test | treatment: F (1, 11) = 35.51, P<0.0001<br>viral expression: F (1, 11) = 7.577, P = 0.0188<br>interaction: F (1, 11) = 10.49, P = 0.0079 |
| 3D | 6 mCherry, 7 hM4 | Mixed-effects model (REML)<br>Factor 1: time (post CNO)<br>Factor 2: viral expression (mcherry, hM4) | time: F (60, 264) = 3.746, P<0.0001<br>viral expression: F (1, 11) = 0.4110, P = 0.5346<br>interaction: F (60, 264) = 0.8477, P < 0.776 |
| 3E | 6 mCherry, 7 hM4 | RM two-way ANOVA<br>Factor 1: treatment (vehicle, CNO)<br>Factor 2: viral expression (mcherry, hM4)<br>Sidak's multiple comparison test | treatment: F (1, 11) = 12.96, P = 0.0042<br>viral expression: F (1, 11) = 0.007926, P = 0.9307<br>interaction: F (1, 11) = 23.64, P = 0.0005 |
| 3F | 6 mCherry, 7 hM4 | Mixed-effects model (REML)<br>Factor 1: time (post CNO)<br>Factor 2: viral expression (mcherry, hM4) | time: F (60, 266) = 2.602, P<0.0001<br>viral expression: F (1, 11) = 39.28, P<0.0001<br>interaction: F (60, 266) = 2.202, P<0.0001 |
| 3G | 6 mCherry, 7 hM4 | RM two-way ANOVA<br>Factor 1: treatment (vehicle, CNO)<br>Factor 2: viral expression (mcherry, hM4)<br>Sidak's multiple comparison test | treatment: F (1, 11) = 76.99, P<0.0001<br>viral expression: F (1, 11) = 8.633, P = 0.0135<br>interaction: F (1, 11) = 29.00, P = 0.0002 |
| 3I | 5 mCherry, 7 hM3 | RM two-way ANOVA<br>Factor 1: treatment (vehicle, CNO)<br>Factor 2: viral expression (mcherry, hM3)<br>Sidak's multiple comparison test | treatment: F (1, 10) = 4.824, P = 0.0528<br>viral expression: F (1, 10) = 6.35, P = 0.0304<br>interaction: F (1, 10) = 5.893, P = 0.0356 |
| 3J | 9 mCherry, 13 hM3 | unpaired t test (two-tailed) | P = 0.0207 |
| 4I | 5 mCherry, 6 hM3 | RM two-way ANOVA<br>Factor 1: treatment (vehicle, CNO)<br>Factor 2: viral expression (mcherry, hM3)<br>Sidak's multiple comparison test | treatment: F (1, 9) = 0.008375, P = 0.9291<br>viral expression: F (1, 9) = 29.24, P = 0.0004<br>interaction: F (1, 19) = 39.34, P = 0.0001 |
| 4J | 5 mCherry, 5 hM3 | unpaired t test (two-tailed) | P<0.0001 |
| 4K | 5 mCherry, 6 hM3 | RM two-way ANOVA<br>Factor 1: time (post CNO)<br>Factor 2: viral expression (mcherry, hM3) | time: F (3.6, 32.4) = 5.038, P = 0.0037<br>viral expression: F (1, 9) = 6.141, P = 0.0351<br>interaction: F (12, 108) = 2.535, P = 0.0055 |
| 4L | 5 mCherry, 6 hM3 | RM two-way ANOVA<br>Factor 1: treatment (vehicle, CNO)<br>Factor 2: viral expression (mcherry, hM3)<br>Sidak's multiple comparison test | treatment: F (1, 9) = 7.013, P = 0.0266<br>viral expression: F (1, 9) = 2.447, P = 0.1522<br>interaction: F (1, 9) = 10.42, P = 0.0104 |
| 4M | 5 mCherry, 6 hM3 | RM two-way ANOVA<br>Factor 1: time (post CNO)<br>Factor 2: viral expression (mcherry, hM3) | time: F (2.547, 22.92) = 2.704, P = 0.0769<br>viral expression: F (1, 9) = 0.2985, P = 0.5981<br>interaction: F (12, 108) = 0.3523, P = 0.9766 |
| 4N | 5 mCherry, 6 hM3 | RM two-way ANOVA<br>Factor 1: treatment (vehicle, CNO)<br>Factor 2: viral expression (mcherry, hM3)<br>Sidak's multiple comparison test | treatment: F (1, 9) = 0.3504, P = 0.5685<br>viral expression: F (1, 9) = 0.03257, P = 0.8608<br>interaction: F (1, 9) = 2.870, P = 0.1245 |
| 4O | 5 mCherry, 6 hM3 | RM two-way ANOVA<br>Factor 1: time<br>Factor 2: treatment/viral expression<br>Sidak's multiple comparison test | time: F (2.319, 41.74) = 966.6, P <0.0001<br>treatment/viral expression: F (3, 18) = 0.6880, P=0.5711<br>interaction: F (12, 72) = 0.8902, P = 0.5606 |

### Supplementary Table 1C: Summary of Statistical Analyses Cont.

| | | | |
|---|---|---|---|
| 5D | 10 mCherry, 9 hM3 | RM two-way ANOVA<br>Factor 1: treatment (vehicle, CNO)<br>Factor 2: viral expression (mcherry, hM3)<br>Sidak's multiple comparison test | treatment: F (1, 17) = 2.219, P = 0.1547<br>viral expression: F (1, 17) = 3.178, P = 0.0925<br>interaction: F (1, 17) = 6.715, P = 0.0190 |
| 5E | 10 mCherry, 9 hM3 | Mixed-effects model (REML)<br>Factor 1: time (post CNO)<br>Factor 2: viral expression (mcherry, hM3) | time: F (38, 253) = 7.182, P< 0.0001<br>viral expression: F (1, 17) = 0.008586, P = 0.9273<br>interaction: F (38, 253) = 3.237, P < 0.0001 |
| 5F | 10 mCherry, 9 hM3 | RM two-way ANOVA<br>Factor 1: treatment (vehicle, CNO)<br>Factor 2: viral expression (mcherry, hM3)<br>Sidak's multiple comparison test | treatment: F (1, 17) = 7.704, P=0.0130<br>viral expression: F (1, 17) = 0.7091, P = 0.4114<br>interaction: F (1, 17) = 3.408, P = 0.0824 |
| 5G | 10 mCherry, 9 hM3 | Mixed-effects model (REML)<br>Factor 1: time (post CNO)<br>Factor 2: viral expression (mcherry, hM3) | time: F (38, 253) = 2.475, P< 0.0001<br>viral expression: F (1, 17) = 5.311, P = 0.0341<br>interaction: F (38, 253) = 1.627, P < 0.0156 |
| 5H | 10 mCherry, 9 hM3 | RM two-way ANOVA<br>Factor 1: treatment (vehicle, CNO)<br>Factor 2: viral expression (mcherry, hM3)<br>Sidak's multiple comparison test | treatment: F (1, 17) = 0.002755, P=0.9588<br>viral expression: F (1, 17) = 1.614, P = 0.2211<br>interaction: F (1, 17) = 17.49, P = 0.0006 |
| 5I | 10 mCherry, 9 hM3 | RM two-way ANOVA<br>Factor 1: time<br>Factor 2: treatment/viral expression<br>Sidak's multiple comparison test | time: F (1.153, 39.19) = 215.6, P <0.0001<br>treatment/viral expression: F (3, 34) = 7.027, P = 0.0008<br>interaction: F (12, 136) = 5.185, P<0.0001 |
| 5M | 3 mCherry, 5 hM3 | RM two-way ANOVA<br>Factor 1: treatment (vehicle, CNO)<br>Factor 2: viral expression (mcherry, hM3)<br>Sidak's multiple comparison test | treatment: F (1, 6) = 1.756, P = 0.2334<br>viral expression: F (1, 6) = 5.031, P = 0.0661<br>interaction: F (1, 6) = 1.004, P = 0.3551 |
| 5N | 3 mCherry, 5 hM3 | RM two-way ANOVA<br>Factor 1: time (post CNO)<br>Factor 2: viral expression (mcherry, hM3) | time: F (4.545, 27.27) = 5.871, P = 0.0011<br>viral expression: F (1, 6) =0.0002876, P = 0.9870<br>interaction: F (27, 162) = 2.267, P = 0.0009 |
| 5O | 3 mCherry, 5 hM3 | RM two-way ANOVA<br>Factor 1: treatment (vehicle, CNO)<br>Factor 2: viral expression (mcherry, hM3)<br>Sidak's multiple comparison test | treatment: F (1, 6) = 0.03797, P = 0.8519<br>viral expression: F (1, 6) = 0.03586, P = 0.8560<br>interaction: F (1, 6) = 0.2161, P = 0.6584 |
| 5P | 3 mCherry, 5 hM3 | RM two-way ANOVA<br>Factor 1: time (post CNO)<br>Factor 2: viral expression (mcherry, hM3) | time: F (2.916, 17.50) = 1.330, P = 0.2965<br>viral expression: F (1, 6) =2.985, P = 0.1348<br>interaction: F (27, 162) = 3.425, P <0.0001 |
| 5Q | 3 mCherry, 5 hM3 | RM two-way ANOVA<br>Factor 1: treatment (vehicle, CNO)<br>Factor 2: viral expression (mcherry, hM3)<br>Sidak's multiple comparison test | treatment: F (1, 6) = 0.2211, P = 0.6548<br>viral expression: F (1, 6) = 0.2390, P = 0.6423<br>interaction: F (1, 6) = 9.596, P = 0.0212 |
| 5R | 3 mCherry, 5 hM3 | RM two-way ANOVA<br>Factor 1: time<br>Factor 2: treatment/viral expression<br>Sidak's multiple comparison test | time: F (2.906, 34.87) = 405.5, P <0.0001<br>treatment/viral expression: F (3, 12) = 28.53, P<0.0001<br>interaction: F (12, 48) = 13.07, P<0.0001 |
| Supplemental 1A | 5 mCherry, 6 hM3 | RM two-way ANOVA<br>Factor 1: treatment (vehicle, CNO)<br>Factor 2: viral expression (mcherry, hM3)<br>Sidak's multiple comparison test | treatment: F (1, 9) = 2.099, P = 0.1813<br>viral expression: F (1, 9) = 0.2308, P = 0.6424<br>interaction: F (1, 9) = 0.7412, P = 0.4116 |

### Supplementary Table 1D: Summary of Statistical Analyses Cont.

| | | | |
|---|---|---|---|
| Supplemental 3A | 5 mCherry, 6 hM3 | RM two-way ANOVA<br>Factor 1: time (post CNO)<br>Factor 2: viral expression (mcherry, hM3) | time: F (4.061, 36.55) = 5.784, P = 0.0010<br>viral expression: F (1, 9) = 0.06638, P = 0.8025<br>interaction: F (12, 108) = 0.3768, P = 0.9692 |
| Supplemental 3B | 5 mCherry, 6 hM3 | RM two-way ANOVA<br>Factor 1: treatment (vehicle, CNO)<br>Factor 2: viral expression (mcherry, hM3)<br>Sidak's multiple comparison test | treatment: F (1, 9) = 1.859, P = 0.2059<br>viral expression: F (1, 9) = 0.4086, P = 0.5386<br>interaction: F (1, 9) = 0.003788, P = 0.9523 |
| Supplemental 3C | 8 mCherry, 9 hM3 | Mixed-effects model (REML)<br>Factor 1: time (post CNO)<br>Factor 2: viral expression (mcherry, hM3) | time: F (38, 225) = 5.370, P <0.0001<br>viral expression: F (1, 15) = 1.478, P = 0.2429<br>interaction: F (38, 225) = 2.233, P = 0.0002 |
| Supplemental 3D | 8 mCherry, 9 hM3 | RM two-way ANOVA<br>Factor 1: treatment (vehicle, CNO)<br>Factor 2: viral expression (mcherry, hM3)<br>Sidak's multiple comparison test | treatment: F (1, 15) = 6.964, P = 0.0186<br>viral expression: F (1, 15) = 2.111, P = 0.1669<br>interaction: F (1, 15) = 0.1738, P = 0.6827 |
| Supplemental 3E | 3 mCherry, 5 hM3 | RM two-way ANOVA<br>Factor 1: time (post CNO)<br>Factor 2: viral expression (mcherry, hM3) | time: F (4.949, 29.69) = 6.543, P = 0.0003<br>viral expression: F (1, 6) = 2.533, P = 0.1626<br>interaction: F (27, 162) =1.874, P = 0.0091 |
| Supplemental 3F | 3 mCherry, 5 hM3 | RM two-way ANOVA<br>Factor 1: treatment (vehicle, CNO)<br>Factor 2: viral expression (mcherry, hM3)<br>Sidak's multiple comparison test | treatment: F (1, 6) = 0.0007374, P = 0.9792<br>viral expression: F (1, 6) = 3.189, P = 0.1244<br>interaction: F (1, 6) = 1.213, P = 0.3129 |
